# GREM1 is epigenetically reprogrammed in muscle cells after exercise training and controls myogenesis and metabolism

**DOI:** 10.1101/2020.02.20.956300

**Authors:** Odile Fabre, Lorenzo Giordani, Alice Parisi, Pattarawan Pattamaprapanont, Danial Ahwazi, Caroline Brun, Imane Chakroun, Anissa Taleb, Alexandre Blais, Emil Andersen, Lars R Ingerslev, Pascal Maire, Michael Rudnicki, Claire Laurens, Kiymet Citirikkaya, Christian Garde, Leonidas Lundell, Atul S Deshmukh, Cédric Moro, Virginie Bourlier, Rémi Mounier, Fabien Le Grand, Romain Barrès

**Author notes:** Equal contribution (Odile Fabre, Lorenzo Giordani, Alice Parisi). Co-last authors (Fabien Le Grand, Romain Barrès). Correspondence: Romain Barrès, Mærsk Tornet 7.7.75, Blegdamsvej 3B, 2200 Copenhagen N, Denmark, mail; Fabien Le Grand, Faculté de Médecine, 8 Avenue Rockefeller, 69008 Lyon, France, mail.

## Abstract

Exercise training improves skeletal muscle function, notably through tissue regeneration by muscle stem cells. Here, we hypothesized that exercise training reprograms the epigenome of muscle cell, which could account for better muscle function. Genome-wide DNA methylation of myotube cultures established from middle-aged obese men before and after endurance exercise training identified a differentially methylated region (DMR) located downstream of *Gremlin 1* (*GREM1*), which was associated with increased *GREM1* expression. GREM1 expression was lower in muscle satellite cells from obese, compared to lean mice, and exercise training restored GREM1 levels to those of control animals. We show that GREM1 regulates muscle differentiation through the negative control of satellite cell self-renewal, and that GREM1 controls muscle lineage commitment and lipid oxidation through the AMPK pathway. Our study identifies novel functions of GREM1 and reveals an epigenetic mechanism by which exercise training reprograms muscle stem cells to improve skeletal muscle function.

## Introduction

Skeletal muscle is a large organ that, in addition to locomotion, accounts for a variety of other physiological roles such as endocrine and metabolic functions^1,2^. Muscle stem cells, *i.e*. satellite cells, continuously regenerate skeletal muscle and participate in muscle tissue adaptation^3,4^, notably in response to physical exercise, where the adaptive response to endurance and resistance exercise is associated with recruitment and differentiation of satellite cells into newly formed myofibers^5–7^.

Several studies showed that satellite cells isolated from a skeletal muscle biopsy and differentiated in culture retain metabolic characteristics of the donor. For instance, cultures derived from satellite cells obtained from type 2 diabetes (T2D) subjects show defects in insulin signaling and glucose^8–12^ and lipid metabolism^13,14^. Similarly, eight weeks of endurance exercise training lead to metabolic adaptations *in vivo* that are partly recapitulated in myotube cultures generated from skeletal muscle biopsies^15^. In particular, biopsy-derived myotube cultures collected post-training showed improved glucose oxidation, glycogen synthesis and glucose-induced inhibition of palmitate oxidation^15^. This indicates that muscle satellite cells are metabolically reprogrammed by endurance exercise training. While genetic factors can cause different phenotypes in satellite cells collected from T2D donors^16,17^, phenotypic changes observed in cells from the same donor after a lifestyle intervention strongly support that epigenetic factors establish a “metabolic memory” after exercise training.

Development of the skeletal muscle cell is under the control of epigenetic factors like DNA methylation^18,19^. Methylation of DNA regulates gene expression by modulating access of the transcription machinery to promoters or by enabling the recruitment of methyl-binding proteins that control chromatin compaction^20^. DNA methylation is a dynamic process that can be influenced by lifestyle factors such as physical activity and nutrition^21,22^. Exercise training can stably modify DNA methylation landscapes and thereby influence gene expression^22–24^, notably by remodeling methyl groups at genes involved in the insulin signaling pathway, muscle energy metabolism and involved in T2D^25^. An intervention as short as three months of knee-extension is associated with DNA methylation changes at genes related to myogenesis, remodeling of muscle architecture and bioenergetics^26^.

In the present study, we aimed to determine if endurance exercise training reprograms the epigenome of muscle stem cells, which could account for altered muscle progenitor function. We profiled whole-genome DNA methylation of myotube cultures established from obese donors before and after endurance exercise training^15^ and identified differential methylation of a cis-regulatory region controlling the expression of Gremlin 1 (*GREM1*). We show endurance exercise training reprograms *GREM1* expression. We provide evidence that *GREM1* controls lineage commitment and energy metabolism through the AMP-activated protein kinase (AMPK) pathway.

## Results

### Endurance exercise training remodels methylation at a genomic region regulating GREM1

To determine if exercise training reprograms the epigenome of muscle satellite cells, we performed methylated DNA capture sequencing in biopsy-derived myotubes described in a previous study^15^. For each subject, satellite cells were collected in a paired fashion: before (pre-training) and after (post-training) an 8-wk endurance exercise-training program. Genome-wide DNA methylation results were plotted on a multidimensional scaling plot with the paired DNA methylation profiles obtained for each subject before and after training. Principal component analysis of DNA methylation levels in both groups shows two well-defined clusters depending on the pre-or post-training condition and highlights a homogeneous response to endurance exercise training in all subjects (Fig. 1a). Analysis of differential methylation returned 116 differentially methylated regions (DMRs) where 102 DMRs were hypomethylated whereas only 14 DMRs were hypermethylated (Fig. 1b). Most DMRs were distant from transcription start sites (at least 50 kb for 60% of the DMRs; Supplementary Fig. 1a), and mainly located in distal intergenic regions and introns (66 and 44 DMRs, respectively; Supplementary Fig. 1.

**Fig. 1.**
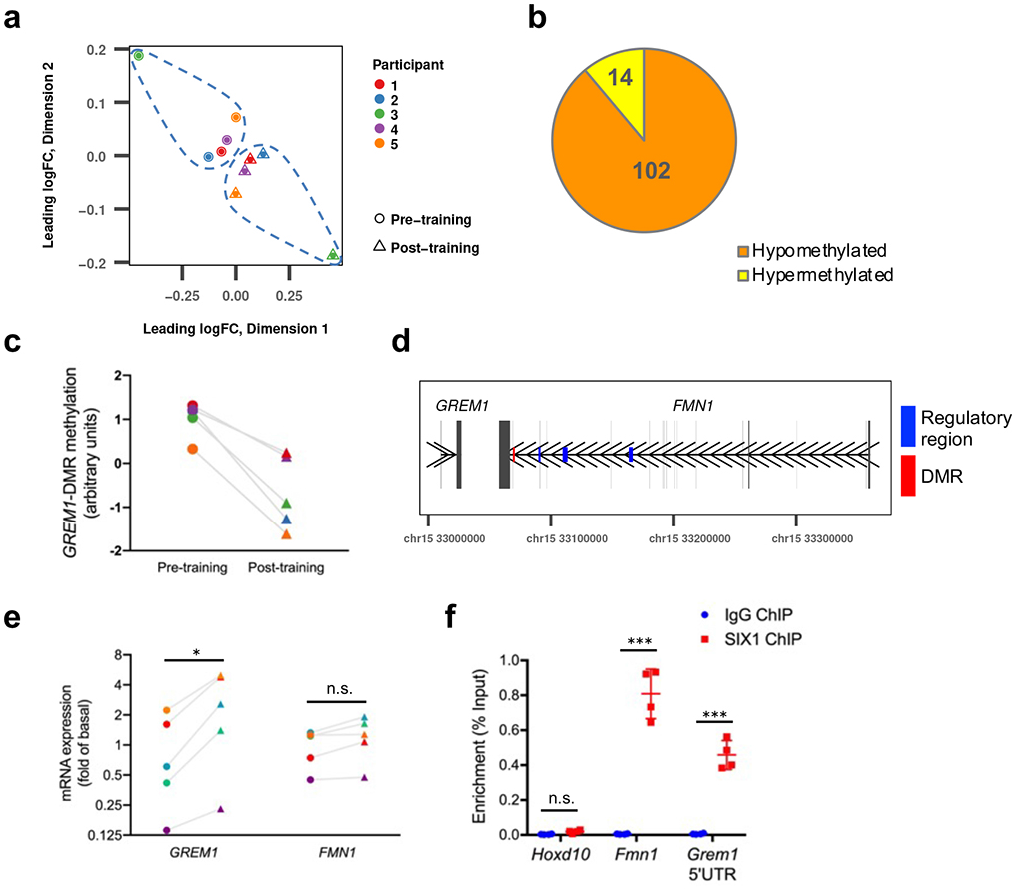
Exercise training stablv remodels DNA methylation in muscle progenitor cells from obese individuals. Muscle cell precursors were isolated from Vastus Lateralis muscle biopsies from five obese individuals before and after exercise training. After differentiation in myotubes, cells were harvested for genomic DNA purification. DNA was then sonicated, and methylated fragments were isolated using methyl-CpG binding domain (MBD)-coupled beads. Captured DNA was used as starting material for preparation of libraries for whole-genome sequencing. n = 5. **a,** Multidimensional scaling plot of methyl-capture sequencing data. **b,** Distribution of identified differentially methylated regions (DMRs) between hypomethylated and hypermethylated. **c,** Methylation levels at the GREM1-DMR. Color codes for participants correspond to those of **a. d,** Localization in chromosome 15q13.3 of GREM1-associated DMR (in red) and of three regulatory regions of Grem1 expression in FMN1 gene (identified in mouse^30^ and converted to corresponding human coordinates, in blue). Each vertical black bar represents an exon. As indicated by the arrows, GREM1 and FMN1 genes are transcribed in opposite orientation. **e,** RT-qPCR analysis of GREM1 and FMN1 mRNA expression in differentiated myotubes from human donors, as means of 2^-ΔCt^ values expressed relatively to the mean of pre-training 2^-ΔCt^ values obtained for the five subjects, which was set at 1. Vertical axis scale is logarithmic base 2. Circle = Pre-training; triangle = Post-training. Color codes for participants correspond to those of **a** and **c. f**, Occupancy of SIX1 at the upstream regions of Grem1 and HoxD10 (control) genes in C2C12 myoblasts. The input represents the relative enrichment of PAX7-Flag proteins compared to the IgG control. Scale bars, 20μm. Error bars indicate SD. *p<0.05; ***p< 0.001; n.s.: not statistically significant.

To investigate possible enrichment for gene ontology terms, we attributed each DMR to the nearest gene based on the shortest distance to the transcription start site. We did not find any enrichment using online Gene Ontology enRIchment anaLysis and visuaLizAtion tool (GOrilla)^27^ or Genomic Regions Enrichment of Annotations Tool (GREAT)^28^. We thus opted for a candidate gene approach and selected *GREM1* based on highest fold change in methylation (3.1-fold less methylated post-training, Fig. 1c). The DMR near *GREM1* is located within the *Formin 1 (FMN1*) gene (Fig. 1d). Transcriptional analysis showed that *GREM1*, but not *FMN1* mRNA was differentially expressed in myotube cultures established from the post-training biopsies (2.8-fold up-regulated, p<0.05) (Fig. 1e). While GREM1 DMR does not overlap with a mapped gene-regulatory region in the human genome, in the mouse genome, a region 9-kb distant to the *Grem1* gene and located within the *Fmn1* locus was identified as a cis-regulatory region for *Grem1*^29,30^. Although the *GREM1* DMR was not found at the exact same distance to the transcription start site in the human genome as compared to the mouse genome, the inverse association between methylation of the *GREM1* DMR and *GREM1* expression strongly supports that *GREM1* DMR is located within a cis-regulatory region for *GREM1*.

In myoblasts from adult mice, *Grem1* regulatory regions were described to contain binding sites for the SIX homeobox 1 (Six1)^31^, a transcription factor which controls satellite cell self-renewal and regeneration of skeletal muscle^32^. Using chromatin immunoprecipitation (ChIP), we confirmed that SIX1 targets the *Grem1* locus in muscle cells. We found SIX1 co-precipitates with the *Grem1* locus, supporting a physical interaction between *Grem1* regulatory regions and SIX1 (Fig. 1f). Our results suggest that reprogramming of *GREM1* expression after exercise training may potentiate satellite cell self-renewal and regeneration, an important function after exercise.

### *GREM1* expression is upregulated upon muscle regeneration and myogenesis

The importance of satellite cells in skeletal muscle regeneration post exercise prompted to investigate the expression pattern of *GREM1* in regenerating adult muscle tissues. We observed a marked induction of *GREM1* expression in *Tibialis Anterior* (*TA*) from wild-type mice during the acute phase of muscle tissue repair induced by cardiotoxin (CTX) injection (Fig. 2a). The surge in *GREM1* expression was similar to the profile of *myosin heavy chain 3* (*Myh3*), a gene coding for the embryonic forms of myosin heavy chains (Fig. 2b).

**Fig. 2.**
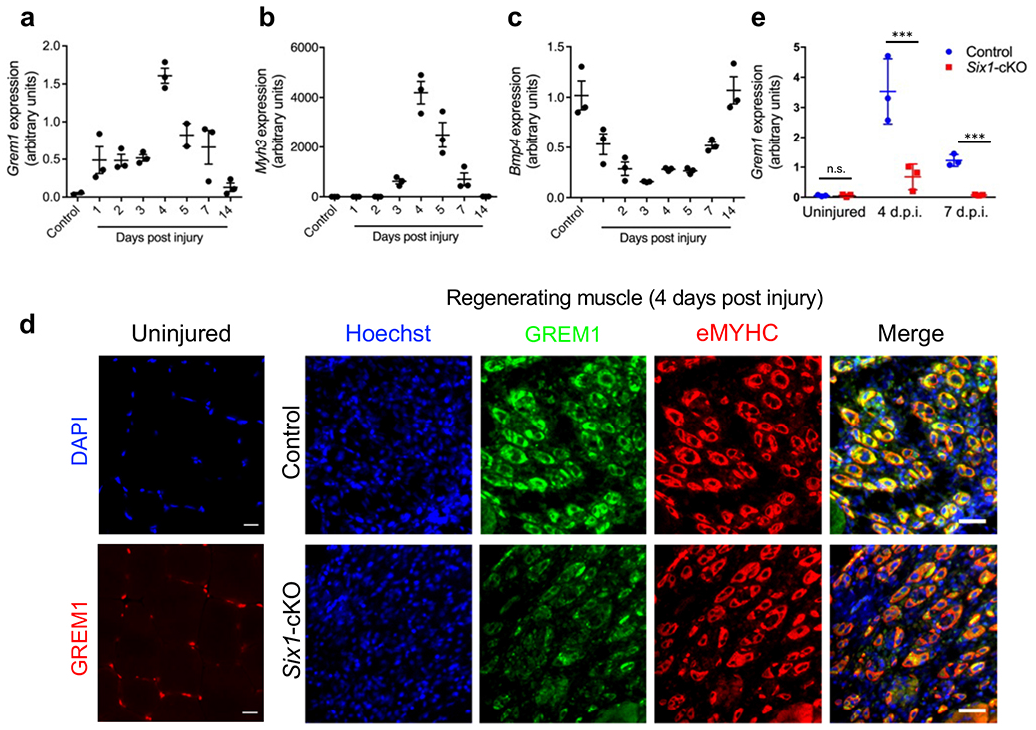
Grem1 is expressed during skeletal muscle regeneration. Gene expression profiling by RT-qPCR during Tibialis Anterior (TA) muscle regeneration for **a**, Grem1, **b**, Myh3, and **c**, Bmp4. **d**, GREM1 (green) immunolocalization on uninjured and regenerating TA muscle cryosections from wild-type and conditional satellite cell-specific Six1 knock-out mice (Six1-cKO). Embryonic MyHC (eMyHC) were stained in red. **e**, RT-qPCR analysis of Grem1 expression in TA muscle of control or Six1-cKO animals. Nuclei were stained with Hoechst or DAPI (blue). Scale bars: 20μm. n = 3 per each experimental time point. Error bars indicate ±SD. ***p<0.001; n.s.: not statistically significant.

GREM1 is an antagonist of the bone morphogenetic protein (BMP) signaling pathway through competitive binding to BMPs^33–35^, as well as inhibition of *Bmp4* transcription^36,37^. Consistent with the literature, *Bmp4* transcripts followed an opposite pattern compared to *GREM1* expression; *Bmp4* expression was low following injury and only returned to basal level once muscle tissue was regenerated, 14 days after injury (Fig. 2c). Immunolocalization of GREM1 protein on muscle tissue cryosections further showed a marked increase in GREM1 expression in newly formed myofibers (Fig. 2d). Interestingly, in mice with targeted *Six1* deletion in muscle satellite cells^32^, Six1 deletion was associated with a low GREM1 expression at both protein and transcript levels after injury (Fig. 2d and e). Thus, our results show that GREM1 expression is induced by muscle regeneration and that such induction is under the control of SIX1.

### GREM1 potentiates myogenesis and antagonizes BMP signaling in primary muscle cells

Along muscle cell differentiation, GREM1 expression at the protein and gene level was markedly increased (Fig. 3a and b) while *Bmp4* expression was gradually decreased (Fig. 3c), which supports our hypothesis that GREM1 plays a role in the differentiation of skeletal muscle cells. To evaluate the effect of *Grem1* on myogenesis, we set primary myoblasts to differentiate in low serum conditions supplemented with either GREM1 or BMP4 and investigated morphological features in the early phases of muscle differentiation. While BMP4 treatment inhibited muscle cell differentiation as previously shown^38,39^ (Fig. 3d-f), GREM1 incubation resulted in the formation of larger myotubes, containing higher number of nuclei (Fig. 3d-f). To investigate the effect of GREM1 on BMP signaling, we measured the activation of BMP signaling by BMP4 following co-incubation by GREM1. We detected that the concomitant addition of GREM1 and BMP4 blocked the effect of BMP4 alone on SMAD phosphorylation (Fig. 3g and h). Moreover, GREM1 blunted the increase in mRNA expression of the BMP target gene *Id1*^40^ (Supplementary Fig. 2). These results demonstrate that GREM1 potentiates myogenesis by antagonizing BMP signaling.

**Fig. 3.**
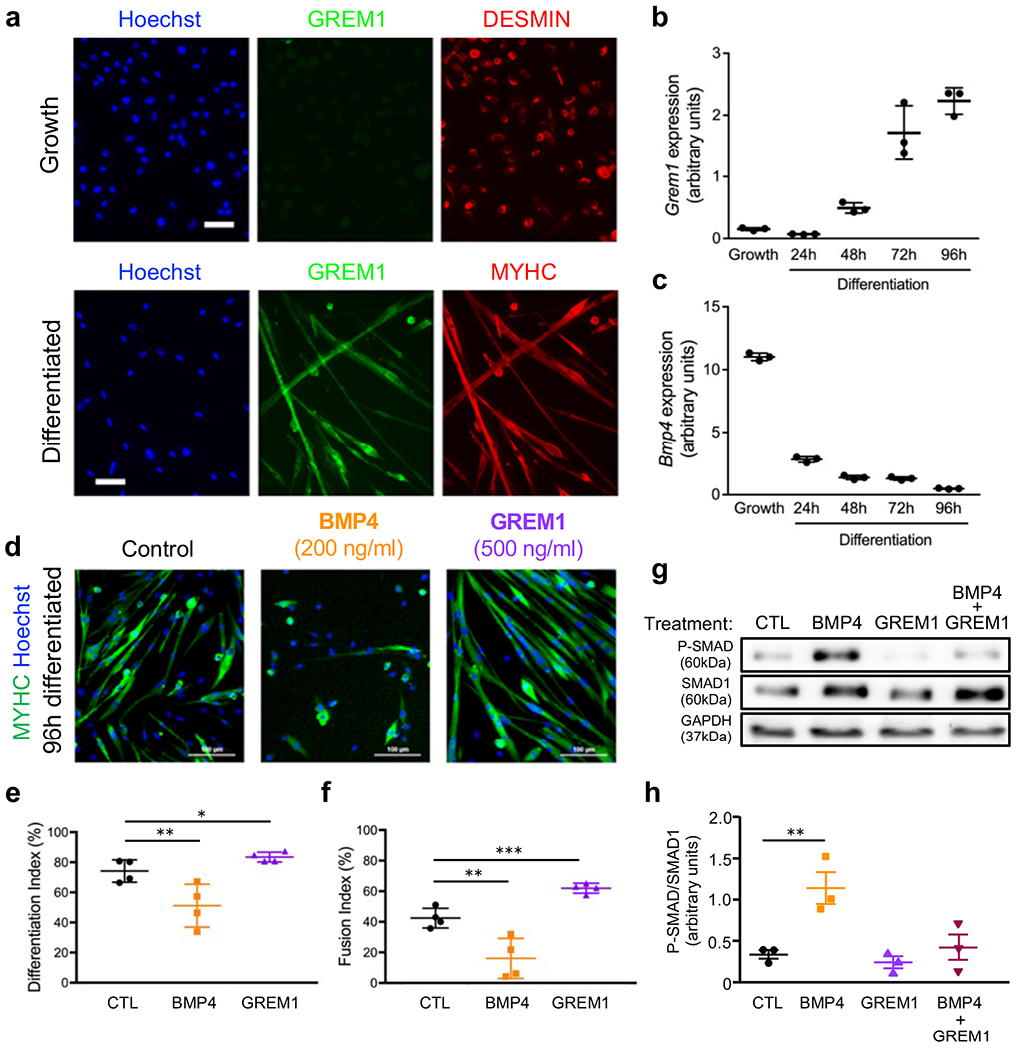
GREM1 antagonizes BMP4 signaling in primary myoblasts. a, Immunolocalization of GREM1 (green) in DESMIN-positive primary myoblasts or in differentiated myotubes expressing MYHC (red). Gene expression profiling by RT-qPCR during in vitro primary myoblast differentiation for b, Grem1 or c, Bmp4. n = 3. d, MYHC immunolocalization (green) of primary myotubes differentiated in presence of GREM1 or BMP4 recombinant proteins. Nuclei were stained with Hoechst (blue). e, Quantification of the differentiation index (percentage of nuclei expressing MYHC). f, Quantification of the fusion index (percentage of nuclei within myotubes). n = 4. g, Western-blot analysis of phosphorylated-SMAD1 proteins in primary myocytes stimulated with recombinant GREM1 or BMP4 or both. GAPDH is a loading control. h, Quantification of phosphorylated SMAD1 over total SMAD1. Scale bars, 20 μm (A), 100 μm (D). n = 3. Error bars indicate ±SD. *p<0.05; **p<0.01; ***p< 0.001; n.s.: not statistically significant.

### GREM1 controls myogenesis through the negative regulation of satellite cell self-renewal

To assess the function of GREM1 on skeletal muscle regeneration *in vivo*, we injured *TA* muscles of adult mice and injected either GREM1 or anti-GREM1 blocking antibodies 3 days after injury (Fig. 4a). Quantification of satellite cell pool by enumeration of the number of sub-laminar Pax7+ cells at day 7 revealed that GREM1 blockade increases satellite cell number under regenerative response (Fig. 4b). While injection of GREM1 did not result in significant changes in myofiber morphology at day 14 (Fig. 4c-d), GREM1 blockade was associated with the formation of smaller myofibers (Fig. 4e) and increased collagen deposition (Fig. 4f), indicating that GREM1 is required for proper muscle tissue regeneration. Quantification of the number of sub-laminar Pax7+ cells at day 14 showed that GREM1 blockade increases the numbers of satellite cells that returned to the niche, while around 50% fewer satellite cells were observed after GREM1 treatment (Fig. 4g). Thus, these data further support that GREM1 controls myogenesis and indicate that GREM1 controls myogenesis through the negative regulation of satellite cell selfrenewal.

**Fig. 4.**
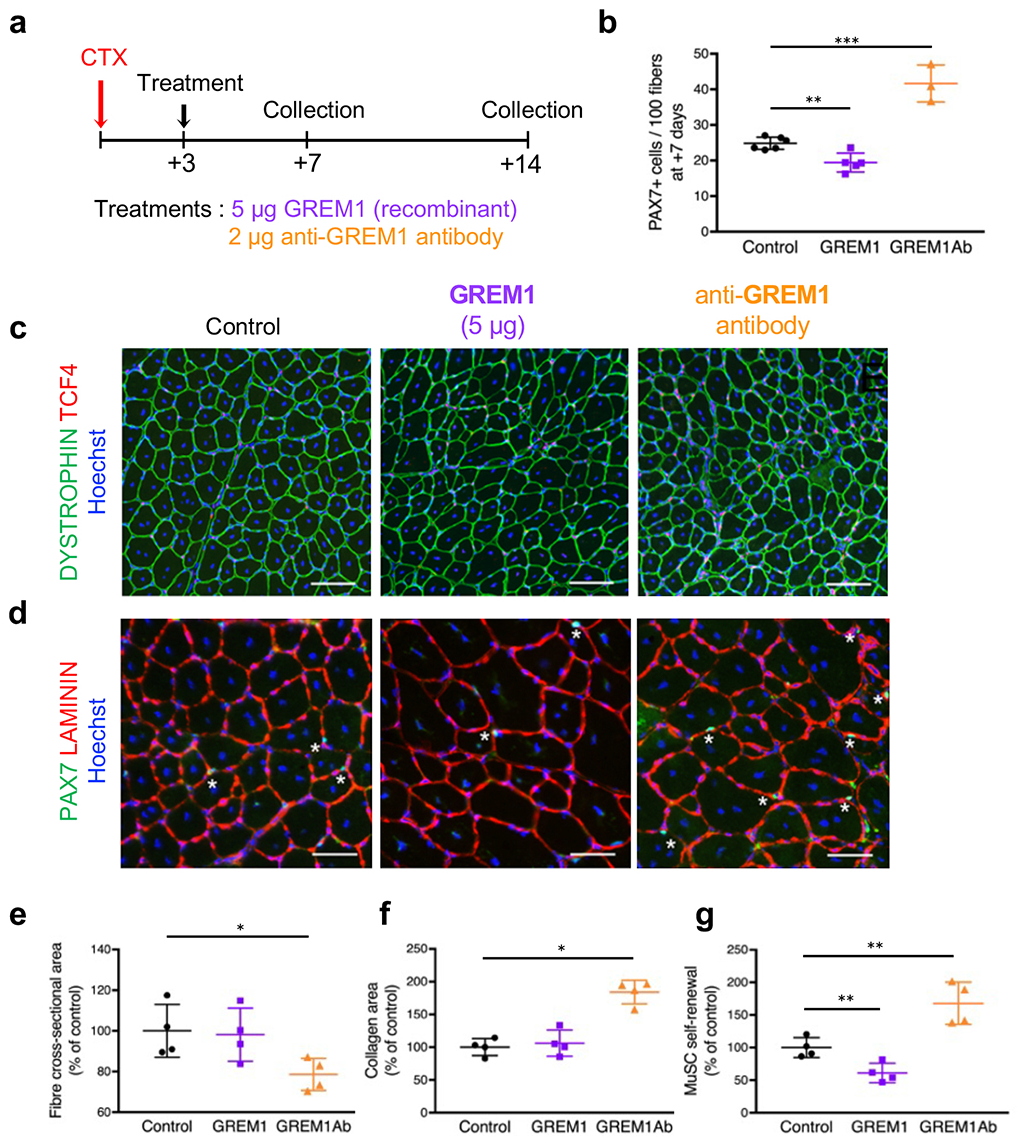
Grem1 limits satellite cell self-renewal in vivo. **a**, Schematic of experimental setup. TA muscles of adult mice were injured by cardiotoxin (CTX) injection, and recombinant GREM1 protein or anti-GREM1 blocking antibodies (GREM1Ab) were administrated 3 days post injury. **b**, Quantification of the number of PAX7+ cells on cryosections generated 7 days after injury in regenerating TA muscles. **c**, Immunolocalization of dystrophin (green) and TCF4 (red) proteins on regenerated TA muscles (14 days post injury). **d**, Immunolocalization of PAX7 (green) and laminin (red) proteins on 14 days post injury in regenerated TA muscles. **e**, Quantification of regenerated myofiber cross-sectional area. **f**, Quantification of area covered by Collagen. **g**, Quantification of muscle satellite cell (MuSCs) self-renewal. Nuclei are stained with Hoechst (blue). Scale bars, 100μm (C), 50μm (D). n = 3 or more for each experimental point. Error bars indicate ±SD. *p<0.05; **p< 0.01; ***p< 0.001. n.s.: not statistically significant.

To investigate if GREM1 impacts self-renewal in a cell-autonomous fashion, we isolated satellite cells by FACS, cultured them for two divisions in the presence of GREM1 or BMP4 proteins (40 hrs, Supplementary Fig. 3a), and quantified the proportion of self-renewing cells (PAX7+/MYOD1+). While BMP4 addition resulted in a higher proportion of self-renewing cells, GREM1 reduced this population and promoted MYOD1 expression by satellite daughter cells (Fig. 5a). Using BrdU assay, we ruled out that the effect of BMP4 or GREM1 is mediated by modulation of cell cycle dynamics (Supplementary Fig. 3b).

**Fig. 5.**
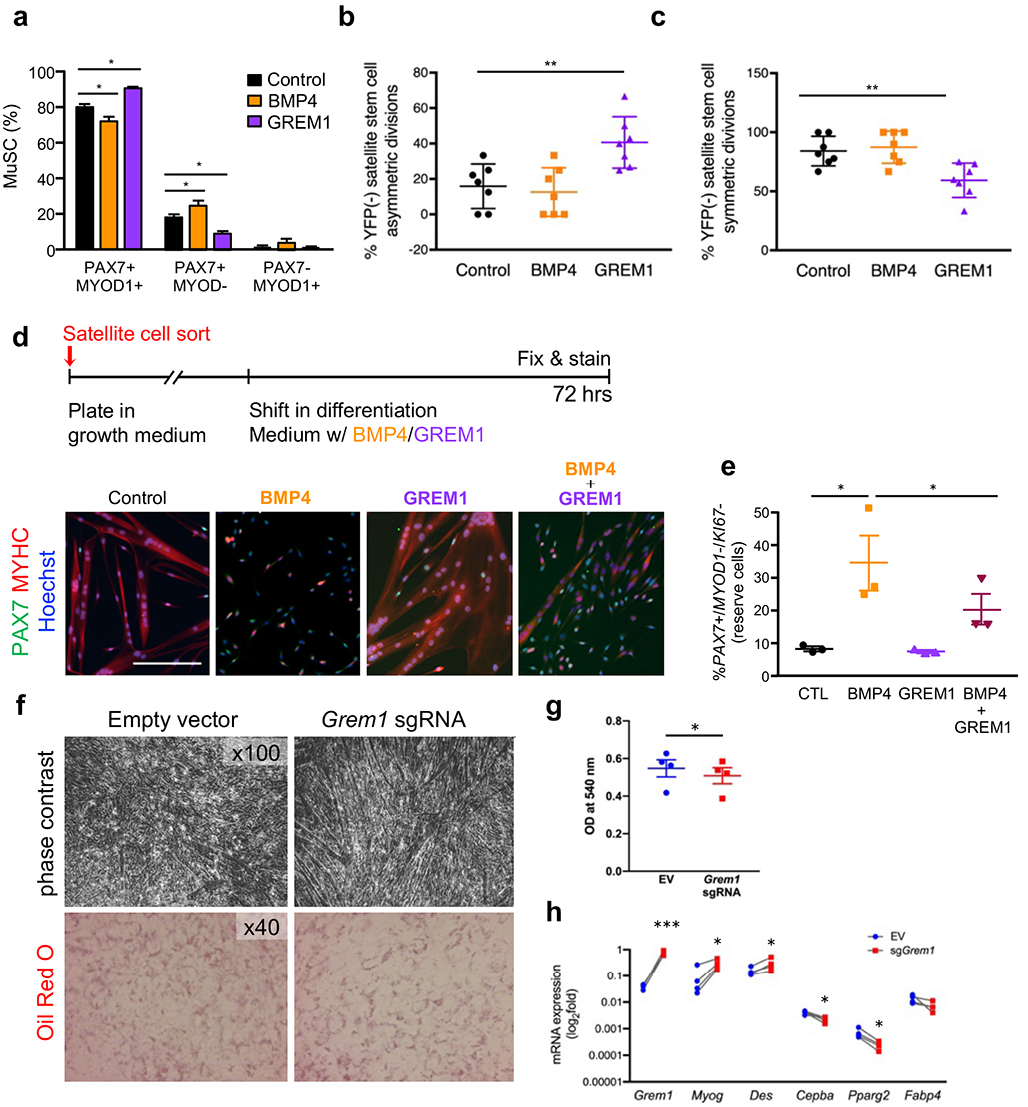
Grem1 promotes satellite cell asymmetric commitment. a, Percentage of muscle satellite cells (MuSC) expressing PAX7 and/or MYOD1. n = 4. b, Quantification of satellite stem cells asymmetric divisions in immunofluorescence images. c, Quantification of satellite stem cells symmetric divisions in immunofluorescence images. n = 7. d, Schematic of experimental setup (top) and representative immunofluorescence image (bottom). Satellite cells were amplified in vitro and induced to differentiate in low mitogen medium containing recombinant GRAM1 and/or BMP4 proteins. Cells were fixed after 72 hrs and stained for PAX7 (green) and MYOD1/MYHC (red) proteins. Nuclei are stained with Hoechst (blue). Scale bar 200 μm. e, Quantification of reserve cells. n = 3. f Representative images of control or GREM1 over-expressing cells under adipogenic differentiation medium. At day 3 of differentiation, representative photomicrographs were taken and then cells were fixed and stained with Oil Red O for intracellular lipid detection. g, Cell-retained Oil Red O was eluted and optical density of the eluates was measured at 540 nm in order to quantify intracellular lipids. n=4. h, Cells were harvested at day 3 of differentiation for total RNA purification and analysis of Grem1, Bmp4, Myog, Des, C/ebpa and Fabp4 mRNA expression by RT-qPCR. n = 4. Error bars indicate SD. *p<0.05; **p<0.01; n.s.: not statistically significant.

Satellite cells consist of a heterogeneous population composed of *Myf5*- lineage marked satellite committed cells and *Myf5*-lineage negative satellite stem cells^41^. Since most of the self-renewal capacity of the satellite cell population is attributable to satellite stem cells^42,43^, we analyzed the impact of BMP4 and GREM1 protein stimulation on satellite stem cell divisions in their niche at the surface of single myofibers prepared from M?5-Cre*ROSA-YFP mice (Supplementary Fig. 3c). We found that GREM1 stimulates asymmetric commitment of satellite stem cells (Fig. 5b) at the expense of symmetric expansion (Fig. 5c). In this experimental setup, BMP4 stimulation did not alter the outcome of satellite stem cell divisions, possibly due to the already high levels of BMP4 expression in these cells^44^. Yet, to validate the effect of BMP4 on promoting satellite cell expansion, we used our *reserve* cell system^45^. We measured the number of satellite cells remaining undifferentiated in differentiation medium and which generates PAX7+/MYOD1’/KI67’ cells at the periphery of differentiated myotubes (Fig. 5d). In this setup, BMP4 stimulation resulted in a striking increase in the *reserve* cells population. This effect was blunted by the addition of GREM1 (Fig. 5e). Altogether, our results indicate that GREM1 is a negative regulator of satellite cell self-renewal by promoting satellite stem cell asymmetric division.

### *Grem1* activates muscle cell differentiation under adipogenic conditions

Since our observation of reprogrammed Grem1 expression was made in obese individuals, and given the well-described function of GREM1 in the inhibition of the BMP pathway during adipogenesis^46^ we tested the hypothesis *Grem1* plays a prominent role in myogenesis under an adipogenic milieu. To perform these experiments, we first generated a model of stable over-expression of endogenous *Grem1* in mouse myogenic C2C12 cell line using the CRISPR/Cas9 Synergistic Activation Mediator-based technique^47^, and confirmed that *Grem1* overexpression was not associated with up-regulation of the predicted off-target promoters of the sgRNA used (Supplementary Fig. 4a). We also checked that *Grem1* over-expression did not affect cell viability (Supplementary Fig. 4b), and that *Grem1* expression was stable across cell division (Fig. 4c). Consistent with the described paracrine function of GREM1^37^, we detected higher extracellular levels of GREM1 in sgRNA-transfected cells (Supplementary Fig. 4d). Overexpression of endogenous GREM1 in adipogenic conditions resulted in higher myotube formation in the late stage of differentiation (Fig. 5f). In addition, we detected fewer cells containing lipid-droplets as showed by Oil-Red O staining and quantification (Fig. 5f and g). Levels of myogenic markers *Des* and *Myog* were higher (*P*=0.016 and *P*=0.021, respectively), while the adipogenic markers CCAAT/enhancer binding protein alpha (*C/ebpa*) and peroxisome proliferator-activated receptor (PPAR) gamma 2 (*PpargC*) were reduced under adipogenic milieu (p=0.035 and p=0.002, respectively; Figure 5H). Collectively, these results show that *Grem1* is improving commitment to myogenesis under adipogenic condition, which may be relevant in the context of obesity *in vivo*.

### Endurance exercise training normalizes GREM1 expression in muscle satellite cells from obese mice

Since our primary observation of increased GREM1 expression was made in satellite cells derived from obese individuals after training, we sought to measure GREM1 *in vivo* expression levels in satellite cells of skeletal muscle from lean and obese mice, before and after training. We found higher GREM1 levels in *soleus* from mice after training, and a trend in the *Plantaris* (*p*=0.06), while GREM1 levels were lower in *ExtensorDigitorum Longus (EDL*) (Fig. 6a and b). At the untrained state, we found GREM1 levels in *plantaris* and *TA* were lower in obese ob/ob mice compared to lean, wild-type animals (Fig. 6b). Interestingly, GREM1 levels in ob/ob mice were normalized by exercise training to lean control levels in TA, *Soleus, EDL*, and with a trend for *plantaris* (*p*=0.11; Fig. 6b). Thus, these results support that GREM1 is down-regulated in obesity and that endurance exercise training increases expression of GREM1 in satellite cells *in vivo*.

**Fig. 6.**
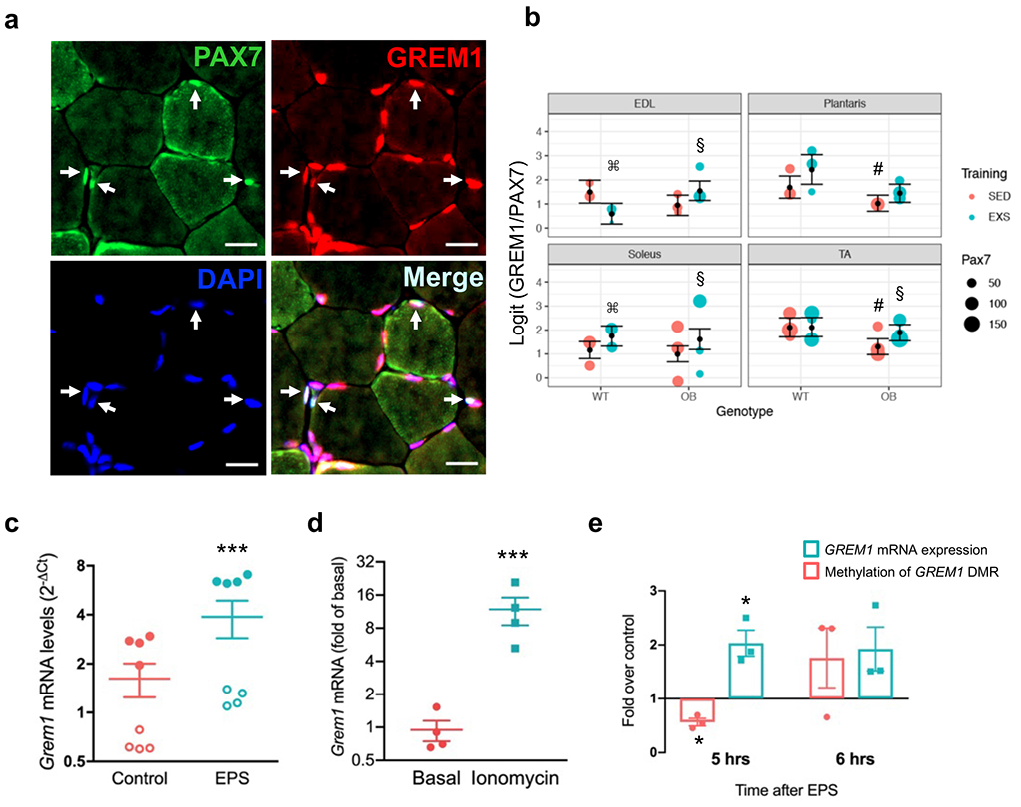
Exercise induces Grem1 expression in vivo and after EPS contraction in myotubes. **a**, Example of immunofluorescence staining images in muscle from mice subjected to 4 wks of exercise training. PAX7 (green), GREM1 (red), DAPI (blue) and colocalization (light blue). Scale bar 20 μm. **b**, Quantification of GREM1-expressing satellite cells in EDL, Extensor Digitorum Longus (EDL), Plantaris, Soleus and Tibialis Anterior (TA). SE = sedentary (red) EXS = exercise-trained (blue). n = 3 animals per group. The number of PAX7+ cells counted under each condition is represented by dot size. ⌘ means p<0.05 for effect of exercise training in wild-type animals; # meansp<0.05 for effect of genotype in sedentary; § meansp<0.05 for effect of exercise training in ob/ob mice. c, Electrical pulse stimulation (EPS) induces GREM1 expression. GREM1 mRNA levels were measured by RT-qPCR in human primary myotubes in the rested state (control) or after 24 hrs of EPS. ***p<0.001. Vertical axis scale is logarithmic base 2. **d**, Acute ionomycin exposure induces Grem1 expression. At day 3 of differentiation, myotubes were treated with 1 μM ionomycin for 3 hrs after which they were harvested for total RNA purification and analysis of Grem1 mRNA expression by RT-qPCR. Each individual dot in the graphs represents the mean of 2^-ΛCt^ duplicate (nontreated cells) or triplicate (treated cells) values from an independent experiment. Vertical axis scale is logarithmic base 10. **e**, Grem1 mRNA expression (blue) and methylation levels of Grem1-DMR (red) at 5 and 6 hrs after EPS stimulation. Differentiated C2C12 cells were subjected twice to EPS at 30 V, 2 ms, 1 Hz for 30 min. Grem1 mRNA expression was detected by RT-qPCR and GREM1 DMR methylation levels were assayed by DNA methylation capture-PCR. Error bars indicate ±SEM in (B) and (E), and ±SD in (C) and (D); *p<0.05; ***p<0.001.

### Muscle cell contraction acutely induces *Grem1* hypomethylation and concomitant mRNA expression

Gene expression response to exercise in skeletal muscle is mediated through several factors including muscle contraction and hormonal input, which drive intracellular cascades leading to gene activation or repression of exercise-responsive genes^48^. To investigate if muscle contraction is sufficient to induce *Grem1* expression, we used electrical pulse stimulation (EPS) to stimulate myotubes in culture and analyzed *Grem1* expression by RT-qPCR following 24hrs of EPS-mediated contraction. We observed a 1.7 to 3.6-fold increase of *Grem1* mRNA level in stimulated human myotubes (p<0.001; Fig. 6c). Increased intracellular calcium levels may represent an intracellular mediator of *Grem1* increased expression after contraction. We next tested the effect of the calcium releasing compound ionomycin^49–51^ on *Grem1* expression in myotubes. *Grem1* mRNA expression was markedly increased by ionomycin (*p*<0.001; Fig. 6d). To determine if EPS-induced *Grem1* expression is associated with DNA methylation remodeling at the *cis*-regulatory region downstream of *Grem1*, we performed RT-qPCR and DNA methylation capture followed by targeted qPCR in a time-course fashion. We found that *Grem1* regulatory region is markedly hypomethylated 5 hrs after EPS and that DNA methylation remodeling is associated with increased mRNA expression (Fig. 6e). Thus, contraction is sufficient to induce *Grem1* expression and support that exercise-induced DNA methylation change of *Grem1* regulatory region controls *Grem1* expression.

### *Grem1* potentiates lipid metabolism in muscle cells

To gain more insight into the physiological role of GREM1, we analyzed *Grem1* expression in muscle cells derived from obese subjects and various clinical parameters of the donor. Interestingly, *Grem1* expression was positively associated with circulating adiponectin levels (*p*=0.010), and was negatively associated with the Insulin Sensitivity Index (ISI) of the liver (*p*=0.013) and blood insulin during the oral glucose tolerance test (*p*=0.023) (Supplementary Fig. 5). These correlations prompted us to hypothesize that *Grem1* participates in the enhancement in energy metabolism as we previously reported in cultures established from these donors^15,52^. Using our CRISPR/Cas9 Synergistic Activation Mediator-based overexpression of endogenous GREM1, we found that while basal glucose uptake, glucose oxidation, glycogen synthesis (Fig. 7a) and insulin-stimulated AKT phosphorylation (Supplementary Figure 5b) were not affected, palmitate oxidation was increased in *Grem1* over-expressing myotubes for both palmitate concentrations tested, regardless of the presence of glucose in the medium (Fig. 7c). These results show that GREM1 improves lipid utilization.

**Fig. 7.**
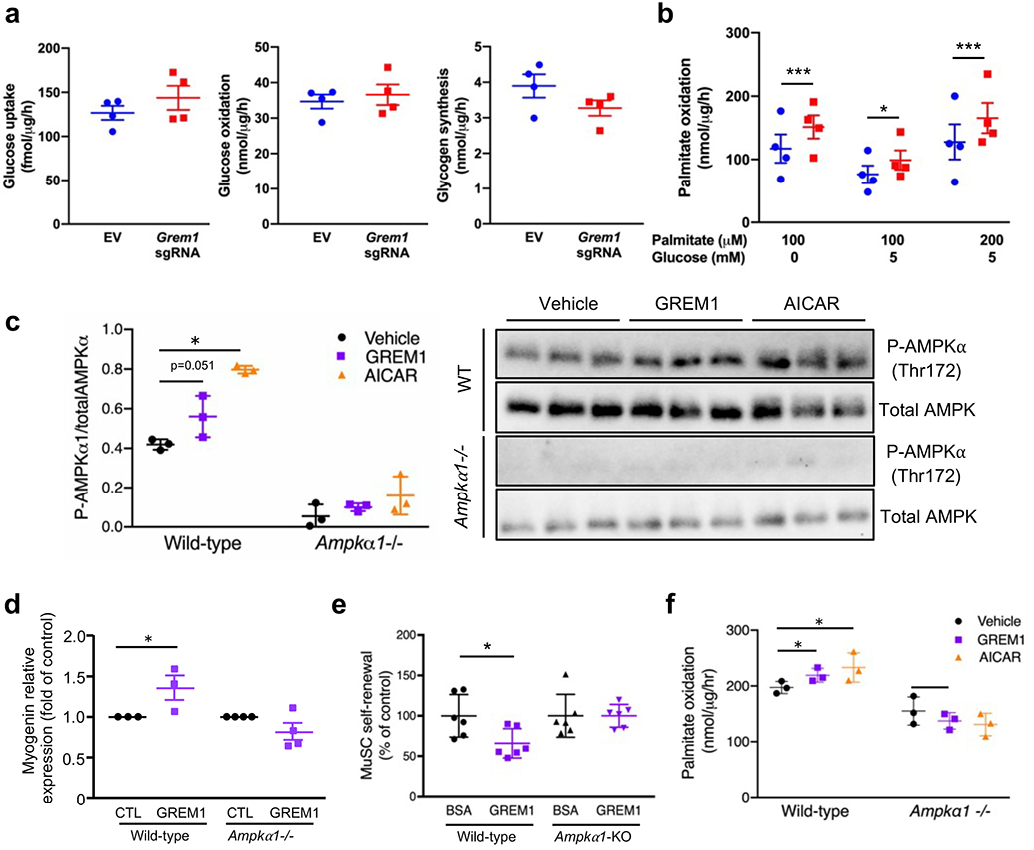
Grem1 increases muscle cell lineage commitment and fatty acid oxidation in myotubes through the AMPK pathway. Before reaching confluence, C2C12 MS2-CAS cells were transfected with Grem1 sgRNA or empty vector for 24 hrs and switched to myogenic differentiation medium. **a**, After 2-hrs serum and glucose starvation, myotubes were incubated for 10 min with cold and radiolabeled deoxyglucose at 100 μM for glucose uptake assay or for 3 hrs with cold and radiolabeled glucose at 5 mM for glucose oxidation and glycogen synthesis assays. n = 4. **b**, After 2 hrs serum starvation, myotubes were incubated for 3 hrs with cold and radiolabeled palmitate at 100 ¡μM or 200 μM in presence or not of 5 mM glucose for assessment of%palmitate oxidation. The graph represents mean DPM values normalized with cell protein content. n = 4. **c**, Phosphorylation of the alpha subunit of%AMPK was assessed (Thr172) in wild-type and Ampka1-nullprimary myoblasts incubated with recombinant GREM1 (500 ng/mL) or AICAR (0.5 mM) for 24 hrs. Western-blot images and the phospho-AMPK to total AMPK ratio was quantified and plotted. n = 3. **d**, RT-qPCR analysis of myogenin expression in wild-type and Ampka1-null primary myoblasts (Ampkod^-/^’) stimulated with recombinant GREM1. n = 3. e, Quantification of the relative amount of sub-laminar PAX7-expressing satellite cells (MuSCs) on muscle cryosections at 14 dpi. in conditional satellite cell-specific Ampka1 knock-out mice (Ampka1-KO). n = 3. f, Palmitate oxidation in fully differentiated myotubes from wild-type or Ampkα1^-/-^-specific cells treated with recombinant GREM1 or AICAR for 3 hrs. n = 3. Error bars indicate ±SD. p<0.05; ***p<0.001; n.s.: not statistically significant.

### AMPK signaling mediates *Grem1* action on satellite cell self-renewal and energy metabolism

To identify the intracellular pathway mediating GREM1 action, we performed proteomic and transcriptomic analyses of serum-starved myoblasts, stimulated with GREM1 or BMP4. We observed that a substantial proportion of modulated transcripts were common between the two conditions (Supplementary Fig. 6a). As expected, GREM1 increased the expression of genes related to muscle cell differentiation, while BMP4 stimulated the expression of genes involved in proliferation (Supplementary Fig. 6b). Gene ontology analysis of GREM1-upregulated genes confirmed the high prevalence of annotations associated with muscle tissue development and function (Supplementary Fig. 6c). Interestingly, Ingenuity Pathway Analysis revealed that GREM1 regulates the protein kinase A (PKA) signaling pathway (Supplementary Fig. 6d), among which many genes related to 5’AMP-activated protein kinase (AMPK), a master regulator of energy metabolism which was recently implicated in satellite cell self-renewal^53^. Using RT-qPCR, we validated that *protein kinase AMP-activated catalytic subunits alpha-1, alpha-2 and gamma-3 (Prkaa1, Prkaa2* and *Prkag3*) are modulated by GREM1 stimulation (Supplementary Figure 6e). Proteomic analysis of myotubes overexpressing endogenous *Grem1* showed that the α1 subunit of the AMPK complex is altered by GREM1 overexpression. Altogether, these results confirmed the importance of GREM1 in skeletal muscle development and prompted us to further investigate the contribution of AMPKα1 on *Grem1* effects. Using primary myoblasts from wildtype and muscle specific *Ampkα1*-null mice^53^, we found that recombinant GREM1 stimulation for 24 hrs trended to increase AMPKα phosphorylation (*p*=0.051; Fig. 7c), and was abolished in *Ampkα1* deficient mice (Fig. 7c). We also found that GREM1 failed to induce myogenin expression in absence of *Ampkα1* (g. 7). *In vivo*, the effect of GREM1 injection on muscle satellite cell self-renewal in *TA* muscle after injury was blunted in *Ampkα1*-KO where the knock-out in satellite cells was induced by tamoxifen, compared to mock-injected controls still expressing *Ampkα1* (ig. 7). Most interestingly, incubation of primary myoblasts from the *Ampkα1*-null mice with GREM1 failed to increase palmitate oxidation, as compared with cells from wild-type animals (Figure 7F). Thus, our data show that GREM1 controls satellite cell self-renewal and lipid oxidation through AMPK signaling.

## Discussion

Here, we characterized the DNA methylation signature of cell cultures derived from skeletal muscle biopsies collected before and after endurance training. We identified that exercise training epigenetically reprograms *Grem1* expression in these cells. Incubation of muscle cells with recombinant GREM1 or over-expression of the endogenous gene highlighted the role of GREM1 in controlling satellite cell self-renewal, muscle cell differentiation and energy metabolism. We identified both GREM1 action on satellite cell homeostasis and GREM1-activated lipid metabolism in muscle cells are controlled by the AMPK pathway. Our results identify a novel function of GREM1 in skeletal muscle biology and support that epigenetic remodeling of *Grem1* participates in an altered muscle stem cell function after endurance training.

Several research groups have investigated genome-wide DNA methylation in human skeletal muscle after exercise training^54^ Differential methylation after exercise was reported after 7-week resistance training^55^, after 6-month endurance training^25^, after 3-month one-legged endurance training^26^, or 3 months of endurance, resistance or combined training^56^. Consistent with our results in muscle cells derived from endurance trained individuals, some studies reported more regions hypomethylated than hypermethylated after training^25,55^. Since these studies were conducted using either resistance^25^ or endurance exercise^55^, the nature of exercise does not to be at play in the overall hypomethylation pattern. We previously reported that acute exercise was associated with hypomethylation at promoter of genes related to energy metabolism and mitochondrial function in skeletal muscle^21^. Since hypomethylation is classically described to be associated with increased transcription, these data suggested that DNA methylation after exercise allows for increased mRNA expression. Overall hypomethylation as a way to enable gene expression is consistent with the global increase in gene expression reported in human skeletal muscle after exercise^56,57^. Our methylated DNA sequencing analysis identified a DMR localized in the intron of *FMN1* gene but closer to the transcription start site of the *Grem1* gene. This suggested that the DMR is located in a regulatory region of *GREM1*. Several instances exist of intronic regions acting as *cis*-regulatory elements for nearby genes, notably Fat mass and obesity-associated (*FTO*) gene where an intronic element regulates expression of the nearby gene Iroquois homeobox 3 (*IRX3*) by establishing long-range interactions^58^. The existence of cis-regulatory elements regulating *Grem1* expression but located within the *Fmn1* gene was previously suggested in mouse, based on the phenotypic observation that the abnormal skeletal pattern in both *Grem1* mutants and mutants of the nearest gene *Fmn1* are similar^59^. Following to these observations, a transcriptional enhancer for *Grem1* located within the 3’ end of the *Fmn1* locus was further identified^30^ and functionally validated in a spontaneously arising deletion of the *Fmn1* coding region^29^. Although the DMR that we identified in the human genome is located at distance of the aforementioned regulatory region identified in the mouse genome, the common regulatory function of these regions on GREM1 expression suggests that the human region is syntenic to the mouse region.

GREM1 is a secreted protein that binds to the BMPs and prevents interaction between BMPs and their receptors^33–35,60^. A role of the balance between GREM1 and BMPs on cell proliferation and differentiation was shown in adipose tissue from obese individuals, where ***Grem1*** and another endogenous BMP inhibitor, *Chordin-like 1* (***CHRDL1***), were increased in concert, thereby inducing resistance to the pro-adipogenic effect of BMP4^46^. In myogenesis too, a role of the *ratio* between BMP4 and GREM1 was suggested^36^. However, in our study, the increase of *Grem1* expression in biopsy-derived myotubes obtained post-training is unlikely to account for a compensatory mechanism targeting the action of BMPs, since neither *BMP4* nor *BMP2* transcript levels were changed in myotubes isolated from post-training biopsies compared to those isolated pre-training (data not shown). A balance between GREM1 and BMPs may not be relevant in isolated muscle cells since in skeletal muscle, GREM1 and BMP4 are expressed by different cell types; while GREM1 is mainly expressed by mature myofibers, BMP4 is exclusively secreted by interstitial cells^36^. Thus, if regulatory mechanisms between GREM1 and BMP4 exist in skeletal muscle, they are more likely to be exerted at the paracrine rather than at the autocrine level.

GREM1 has been studied in numerous cell types where a role on lineage commitment has been reported. In colonocytes, GREM1 participates in acquisition of stem cell properties^61^, while it is involved in kidney formation^62^. Recently, a short-term effect of GREM1 on insulin-signaling in mature human primary myotubes has been identified^63^. Two to four hours incubation of human primary muscle cells with recombinant GREM1 impaired insulin-mediated serine phosphorylation of AKT^63^. In our hands however, overexpression of endogenous GREM1 did not to affect insulin signaling, although we did not test acute effects of GREM1 on insulin signaling. Thus, these differences may be due to the timing in GREM1 stimulation.

Our results demonstrate that GREM1 signals from the regenerating myofibers and inhibit satellite cell self-renewal and promotes commitment. The main role of satellite cells during tissue regeneration is to generate differentiated progeny that can fuse and give rise to new myofibers^64,65^. Stem cell self-renewal is crucial for the maintenance of the niche and tissue homeostasis. Dysfunction on this system leading to decrease self-renewal would lead to depletion of the stem cell population and inability of the tissue to repair following injury, while increased self-renewal may result in overproduction of stem cells and potentially tumorigenesis^66^. Skeletal muscle satellite cells are endowed with a remarkable capacity for self-renewal, and serial rounds of injury yield efficient muscle regeneration for each regeneration cycle^67,68^. Our *in vivo* gain-and loss-of-function experiments demonstrate that GREM1 is important for muscle regeneration following cardiotoxin treatment, through the negative control of satellite cell self-renewal. Further experiments investigating the capacity of muscle to regenerate under normal physiological conditions will provide more definitive insight into the role of GREM1 on muscle homeostasis.

In the light of the effect of GREM1 in the regulation of myogenesis, it is legitimate to question if the metabolic improvement that we observed in *Grem1* over-expressing myotubes is simply the result of an increased number of fully differentiated muscle cells. Yet, our observations do not favor this scenario, as incubation of fully differentiated myotubes with recombinant GREM1 for only 3 hrs (a time that would not allow further differentiation of resident myoblasts) was enough to induce palmitate oxidation to levels similar to cells overexpressing endogenous GREM1, thereby supporting that GREM1-mediated increase in energy metabolism is not caused by differences in muscle differentiation.

In addition to the effect of recombinant GREM1 on terminal differentiation under normal cell culture conditions, we observed that the myogenic potential of myoblasts under an adipogenic milieu was more elevated in *Grem1* overexpressing cells. In cells exposed to an adipogenic medium, *Grem1* overexpressing cells was associated with lower accumulation of lipids and higher capacity to differentiate into myotubes. Since adipogenic conditions activate the BMP pathway^46^, our observed myogenic differentiation supports a mechanism by which *Grem1* may play a particular role in muscle differentiation under conditions where the BMP pathway is activated. Thus, training-induced reprogramming of *Grem1* expression in cells from obese subjects may lead to a better “resistance” to the adipogenic environment of obesity and allow better long-term myogenesis.

In recent years, there has been a growing appreciation that changes in energy balance and metabolic status are critical for stem cell function. We found that expression levels of the cellular energy sensor AMPK were modulated by GREM1. We observed that the effect of GREM1 on satellite cell self-renewal and lipid oxidation are mediated by AMPK. These results are in line with previous studies demonstrating that AMPK controls satellite cell self-renewal^53^. This raises the interesting possibility that during satellite stem cell asymmetric division, metabolic regulators could ensure the faithful segregation of cell fate determinants to the appropriate daughter cells.

Here, we postulated that reprogramming of *Grem1* expression after exercise training participates in the metabolic improvement that we had previously reported in the same cells^15^. However, unlike in the human primary cultures established after training, glucose suppressibility was not increased in *Grem1* overexpressing myotubes. In addition, differentiation at day 4 was not changed in satellite cell cultures derived from our participants^15^. In another study however, primary muscle cells cultures established before and after endurance training showed increased lipid accumulation and oxidation at the post-training stage^69^. We can only speculate that, compared to our experimental systems, the different timing, duration and magnitude of GREM1 expression in primary cells from our participants are playing a role in the discrepancy of phenotypes. In addition, other genes near the other 115 DMRs that we identified may participate in modulating the metabolic and lineage commitment phenotype of primary cells from our donors. Identifying the critical genes reprogrammed after exercise and which modify the phenotype of the differentiated skeletal muscle cell warrants further investigation.

In conclusion, the identification of epigenetic remodeling in skeletal muscle cells after endurance training advances our understanding of the cell memory of exercise training. Given the power of exercise training to prevent or treat chronic diseases, a better understanding of the mechanisms by which lifestyle factors improve skeletal muscle function will provide an important opportunity to develop strategies to mimic some of the beneficial effects of exercise training on human health.

## Methods

### Exercise intervention and isolation of primary muscle cells

Subjects’ characteristics and the protocol of the training program were previously described^15^. Briefly, five healthy (34.8 ± 1.5 years old) obese men (body mass index of 32.8 ± 1.1) with low level of physical activity and no family history of diabetes were subjected to a 8-wk exercise training protocol consisting in five sessions *per* week of aerobic exercise performed at a target heart rate corresponding to 35-85% of their VO_2_ max. Paired biopsies of *Vastus Lateralis* skeletal muscle were collected at rest before and after training, using the Bergstrom needle technique^70^. Biopsy-derived primary cell cultures were obtained as previously described^15^. The research was conducted according to the Declaration of Helsinki, and the protocol was approved by the Ethics Committee of Toulouse University Hospitals (no. NCT01083329 and EudraCT-2009-012124-85). Written informed consent was obtained from all participants.

### Nucleic acid purification

Genomic DNA and total RNA were purified from differentiated myotubes with the AllPrep DNA/RNA/miRNA Universal Kit (Qiagen). The quality of recovered DNA and RNA was assessed by the 260/280 *ratio*, determined by spectrophotometry with a NanoDrop 2000 (Thermo Fisher Scientific), as well as using the Agilent RNA 6000 Nano Kit (Agilent Technologies) for RNA. DNA and RNA concentrations were determined with the Qubit dsDNA HS Assay Kit (Life Technologies) and by spectrophotometry with a NanoDrop, respectively. After purification, DNA and RNA samples were stored at −80°C until further analysis.

### Cell differentiation

Myogenic differentiation medium was high-glucose DMEM/Pen Strep 1% supplemented with horse serum 2% (V/V). For adipogenic differentiation, cells were transfected as described above when 50% confluent and the transfection medium was replaced by growth medium supplemented with human insulin 10 μg/ml, 3-isobutyl-1-methylxanthine 0.5 mM, dexamethasone 1 μM and rosiglitazone 1 μM. All reagents for cell differentiation were purchased from Sigma-Aldrich. Fresh differentiation medium was added to cultured cells daily.

### Electrical pulse stimulation (EPS) of primary human cells

Satellite cells from *Rectus Abdominis* of healthy male subjects were isolated and grown in low-glucose supplemented with FBS 10% and various factors (human epidermal growth factor, BSA, dexamethasone, gentamycin, fungizone, fetuin) as previously described^71^. Cells from four independent donors were grown at 37°C in a humidified atmosphere of 5% CO_2_. Differentiation of myoblasts (*i.e*. activated satellite cells) into myotubes was initiated at ~80-90% confluence, by switching to α-Minimum Essential Medium with Pen Strep 2%, and supplemented with FBS 2%, and fetuin (0.5 mg/ml). The medium was changed every other day and cells were grown up to 5 days. Myotubes were submitted to EPS [2 ms pulses, 10 V, 0.1 Hz frequency] for 24 hrs to recapitulate chronic adaptations of exercise training.

### Immunochemistry

After transfection with Grem1 sgRNA or empty vector, C2C12 MS2-CAS were grown and stained directly in plates, following the protocol provided by Cell Signaling Technology for immunostaining of cultured cells. Cells were fixed with 4% formaldehyde (Sigma-Aldrich) for 15 min at room temperature and then incubated in PBS (Gibco)/FBS 5%/Triton X-100 (Sigma-Aldrich) 0.3% for 1 hr. Staining was done by incubating cells with Desmin rabbit monoclonal antibody (Cell Signaling Technology) and then with Texas Red-conjugated goat anti-rabbit IgG antibody (Invitrogen), diluted at 1:100 and 1:500 respectively in PBS/BSA (Sigma-Aldrich) 1%/Triton X-100 0.3%, for 1 hr at room temperature. A coverslip was mounted on cells using Mowiol (Sigma-Aldrich) as a mounting medium, supplemented with DAPI (Cell Signaling Technology) at 1:2000. Stained cells were visualized using a Zeiss Widefield microscope equipped with an Axiocam 506 mono – 6Mpx detector and analyzed with the Zeiss Zen Blue 2012 software. For pictures, the area was chosen based on nuclei staining.

### Reverse transcription and quantitative PCR (RT-qPCR)

Total RNA was purified with TRIzol (Invitrogen) and used as a template for cDNA synthesis using the iScript cDNA Synthesis Kit (Bio-Rad). mRNA expression was then quantified from generated cDNA by qPCR, using Brilliant III Ultra-Fast SYBR Green QPCR Master Mix (Agilent Technologies) on a CFX96 Real-Time System Thermal Cycler (Bio-Rad). Human and mouse primers were designed using Geneious 7.1.7 computer software. qPCR data were analyzed according to the comparative cycle threshold method and were normalized using the GAPDH gene.

### Protein extraction

Three days after induction of differentiation, transfected C2C12 MS2-CAS myotubes were lysed in ice-cold lysis buffer (HEPES 50 mM [pH 7.4], glycerol 10%, IGEPAL 1%, NaCl 150 mM, NaF 10 mM, EDTA 1 mM, EGTA 1 mM, sodium pyrophosphate 20 mM, sodium orthovanadate 2 mM, sodium-pyrophosphate 1 mM and Sigma FAS Protease Inhibitor Cocktail) and cell media were collected for protein precipitation. One ml of cell medium was incubated with 1 ml trichloroacetic acid 20% on ice for 30 min. Samples were centrifuged 5 min at 10000 x g and the pellet was washed twice with 500 μl acetone. The pellets were then air-dried and resuspended in ice-cold lysis buffer. All the chemicals were from Sigma-Aldrich.

### Methylated DNA capture sequencing

Genomic DNA samples were sheared with a Bioruptor Plus sonication device (Diagenode) and cytosine-methylated DNA was isolated using the MethylMiner Methylated DNA Enrichment Kit (Invitrogen). Captured methylated DNA fractions were then ethanol-precipitated and used for library preparation as previously described^72^. Sequencing was done on a HiSeq 2500, single-read 100-bp read length *per* lane.

### Bioinformatic analysis of methylated DNA capture sequencing

Reads were aligned to hg19 using the Subread aligner^73^, using standard settings except for a consensus threshold of 2 for reporting a hit. Duplicate reads were found and removed using the MarkDuplicates tool of the Picard tools package (http://broadinstitute.github.io/picard). Consensus peaks were found using the Irreproducible Discovery Rate (IDR) framework^74^ with an IDR threshold of 0.005. Reads were counted using the featureCounts tool^75^. Differential peaks were found with edgeR^76^ using settings and methodology described in^77^, analyzing the entire peak. Each peak was then annotated with the nearest gene using ChIPseeker^78^.

### Mice

Mice were handled according to the European Community guidelines. Experimental animal protocols were performed in accordance with the guidelines of the French Veterinary Department and approved by the *Sorbonne Université* Ethical Committee for Animal Experimentation.

Animals were exposed to a standard 12:12 light/dark cycles with normal activity and chow. In this study, 8 to 12-week-old wild-type (WT) mice C57/BL6 mice were used. *Pax7^CreERT2^*;*Six1^lox/lox^* (Six1-cKO) and *Pax7^CreERT2^*;*Ampka1^lox/lox^* (Ampkα1-KO) mice were crossed and maintained in an C57/BL6J background and genotyped by PCR.

### Exercise training experiment in mice and immunofluorescence

#### Animal use

Genetically obese (ob/ob) and lean control littermates (C57/BL6) male mice at 9 weeks old were obtained from Charles River (Italy). Animals were housed in the Department of Experimental Medicine, University of Copenhagen. Animals were maintained at 12:12 h normal-phase dark/light cycle with *ad libitum* access to water and diet throughout the experiment. Mice were divided into sedentary where they maintain their normal activity or subjected to 4 weeks of exercise training. All experiments were performed in accordance with the European Directive 2010/63/EU of the European Parliament and of the Council for the protection of animals used for scientific purposes. Ethical approval was given by the Danish Animal Experiments Inspectorate (#2015-15-0201-00792).

#### Exercise training protocol

Mice were familiarized with treadmill exercise at a speed of 7-10 m/min, 10-15 min/day for 4 days. Then, mice were trained at 7.5% inclination, 30 min/day, 5 days/wk for 1 wk at a speed of 6 and 10 m/min for obese and control mice, respectively. For the other three weeks, mice were run for 60 min/day. The speed was increased to 9 and 13 m/min for the obese and control mice, respectively. Mice were stimulated by a mild electric shock at the back of the treadmill if they refused to run.

#### Sample collection

Sample collection was performed 24 hrs after the last bout of exercise training. Left hindlimb muscles (*TA, EDL, Soleus* and *Plantaris*) were dissected and immediately frozen at their physiological length in Tissue-Tek OCT compound (Cell Path) by isopentane cooled in liquid nitrogen. Then, samples were kept in −80 °C for further analysis. *Immunofluorescence detection* - Cross-sectional sections were cut at 10 μm thickness by cryostat under −20 °C. Sections were fixed in 4% paraformaldehyde for 10 min, washed in PBS, then blocked in 3%BSA and 10% AffinipPure Fab Fragment goat anti-mouse IgG(H+L) in PBST for 60 min. Sections were incubated in human anti-mouse gremlin 1 conjugated to Alexa Fluor 594 and Avain anti-mouse Pax7 conjugated to Alexa Fluor 488 overnight at 4°C. Then, sections were washed in PBS, counterstained with DAPI, and mounted with fluoromount. Immunofluorescences were imaged on Axio Scan.Z1 with 10x magnification.

#### Quantification of Grem1 and Pax7

To quantify numbers of Grem1 in satellite cells, all Pax7+ nuclei and Grem1+/Pax7+ nuclei from entire sections were counted by hand. Then, the ratio of Grem1-expressing cells and the total number of Pax7+ cells were calculated. Within each field, all Grem1+ nuclei were counted, subtracted by Grem1+/Pax7+ nuclei and then divided by the number of muscle fibers. Hierarchical generalized linear model was used to analyze differences in Grem1 nuclei between each group. The ratios of Grem1+ to Pax7+ cells were analyzed using a Generalized Linear Mixed Model (GLMM), with a Binomial error distribution and a log-odds link function. The log-odds ratio model was fitted with training status, genotype and tissue as fixed effects and intercept within animal as a random effect. The model was fitted in R^79^ using the glmer function of the lme4 package^80^. Confidence intervals and p-values were calculated using the emmeans package^81^. Figures were generated using ggplot2^82^.

### Induction of Cre activity and cardiotoxin injury

Intraperitoneal injections of tamoxifen from MP Biomedicals at 5 μl per gram body weight of 20 mg/ml diluted in corn oil were administrated to 8 to 12-week-old mice daily for 4 days prior to injury. Mice were anaesthetized by intraperitoneal injection of ketamin at 0.1mg per gram body weight and Xylazin at 0.01mg per gram body weight diluted in saline solution. After having cleaned the mouse hind legs with alcohol, *tibialis anterior* muscles were injected with 50μl of cardiotoxin solution (Latoxan, 12 μM in saline) using an insulin needle.

### FACS-mediated satellite cell isolation

Satellite cells were obtained from adult hindlimb muscles after enzymatic digestion followed by FACS purification. Briefly, after dissection cells were digested for 1.5 hr in Digestion Solution (2mg/ml Collagenase A; 2.4U/ml Dispase II; 10μg/ml DNAse I; in HBSS. All from Roche). After digestion cells were washed with Wash Buffer (0.2% BSA in HBSS) and filtered twice; once with a 100-μm strainer and once with a 45-μm strainer (Corning Life Science). Cells were then stained with the following antibodies: rat CD31, rat CD45, rat Ly6A (SCA1), rat CD106 (eBioscience) and rat α7 integrin (Ablab) and sorted using a FACS Aria II (BD). Satellite cells were isolated as CD31^neg^, CD45^neg^, Sca ^neg^, α7 integrin^pos^, CD106^pos^.

### Single myofiber isolation

Single myofibers were isolated from the EDL muscle by Collagenase type I digestion and gentle triturating, as previously described^43^. Isolated myofibers were cultured in suspension for up to 2 days in 6-well plates coated with horse serum to prevent fiber attachment. Fibers were incubated in plating medium consisting of 15% Fetal Bovine Serum (Hyclone) and 2 ng/ml bFGF (R&D Systems) in DMEM.

### Satellite cell magnetic isolation and primary myoblasts culture

Skeletal muscles of mice were dissected (quadriceps, *TA, EDL, Gastrocnemius, Soleus, Gluteus*) and transferred to a sterile Petri dish on ice. Muscles were incubated in 1,5U/ml of Collagenase B – 2.4 U/ml of DispaseII – 2M of CaCl2 solution for 45min at 37°C with periodic mechanical digestion. Fetal Bovine Serum was added to stop the digestion. After centrifugation, the pellet was resuspended in growth medium consisting of Ham’s F10 (Life Technologies) with 20% FBS (Eurobio), 1% Pen/Strep (Life Technologies), 2.5 ng/μl basic FGF (R&D Systems). Satellite cells were then purified using MACS cell separation system, according to the manufacturer’s protocol. Satellite cells are then let to proliferate and give rise to primary myoblasts after two passages. If needed, primary myoblasts were differentiated using a differentiation medium composed of DMEM (Gibco) with 2% Horse Serum (Gibco), 1% Pen/Strep (Gibco). Cells were treated with recombinant GREM1 and/or BMP4 proteins (R&D Systems) for a minimum of 24 hrs.

### Muscle histology and immunohistochemistry

Muscles were cut at a thickness of 10 μm with a Leica cryostat. Frozen muscles were stored at −80°C. Sections were fixed with 4% PFA in PBS for 20 min and permeabilized with cooled-methanol. Antigen retrieval was performed with the Antigen Unmasking Solution (Vector) at 95°C for 10 min. Sections were then blocked with 4% BSA, 5% Goat serum in PBS during 3 hrs and incubated with primary antibody overnight at 4°C. Alexa Fluor secondary antibody was incubated on sections 1 hr at room temperature and nuclei were stained with Hoechst. Slides were mounted in fluorescent mounting medium from Dako. Primary antibodies used were against Dystrophin (Leica), embryonic MyHC (Santa Cruz), Grem1 (R&D Systems), Laminin (Santa Cruz), Pax7 (Santa Cruz), Tcf4 (Cell Signaling Tech.).

### Chromatin immunoprecipitation

Examination of anti-Six1 ChIP-seq data obtained in mouse primary myoblasts (to be published elsewhere) revealed the existence of several Six1 binding peaks, notably near the transcription start site of Fmn1 isoform NM_001285459, and at the 3’ UTR of all isoforms. Peaks were visible in both proliferating myoblasts and differentiated myotubes. MEF3 sequences (Six1 binding)^31^, mapped using Cisgenome^83^, were found near the center of most peaks.

Chromatin immunoprecipitation assays (ChIP) were performed to validate binding of Six1 at some of these loci, using the method described in^84^. Twenty-five micrograms of chromatin from C2C12 in proliferation and two micrograms of anti-Six1 (Sigma-Aldrich) or normal rabbit IgG (Jackson ImmunoResearch) were used *per* assay. Real-time PCR was performed using the following primer sequences:

Fmn1_prom_Fwd:CAGGGCAGAGCTACAGCAG;
Fmn1_prom_Rev:GGCAGGTTT CATCTGGAAT G;
Grem1_5UTR_Fwd:TGATCCAACTGCTTCTGTGG;
Grem1_5UTR_Rev:TCCTTGGGATGTTCACTTCTG;
Hoxd10_Fwd:GAGAAATCGGACTCACCTTCC;
Hoxd10_Rev: CACATACCCAGGCAGAACG.

Real-time PCR data were obtained with the SYBR Green method and quantitated using method described in^85^. A one-tailed paired t-test, comparing enrichment at *Fmn1* and at *Hoxd10*, was performed to determine the significance of the ChIP enrichment.

### Immunocytochemistry

Muscle cells or fibers were fixed for 8 min with 4% PFA in PBS then incubated with 5% Goat Serum and 0.2% Triton in PBS for 20 min at room temperature. Cells were incubated with primary antibodies during 1 hr at room temperature, followed by several PBS washes and a 1-hr-incubation with the secondary antibodies. Nuclei were stained with Hoechst. Primary antibodies used were against Desmin (Sigma), pan-Myosin Heavy Chains (DSHB), Pax7 (Santa Cruz), GFP (Thermo Fisher) and MKI67 (Santa Cruz).

For proliferation assay, cells were incubated with BrdU for 40 min before fixing them with 4% PFA for 5 min at room temperature. After a brief wash in PBS, cells were denatured with 2M HCl for 30 minutes at 37°C. To neutralize the acid, 5 min of 6 consecutive washes in PBS were performed. Cells were blocked with 2% Goat Serum, 0.2% Tween 20 PBS for 30 min at 37°C before antibody staining.

### Quantitative real-time PCR (qPCR)

RNA was isolated either by RNeasy Mini Plus Kit (Qiagen) or by TRIzol Reagent followed by a DNAse treatment (Life Technologies). The reverse transcription was performed with 100ng of RNA using the High-Capacity cDNA Reverse Transcription kit (Applied Biosystems). Transcript levels were determined by LightCycler 480 Real-Time PCR System from Roche, using SYBR green I Master from the same company. The melting curves were checked for each experiment and the primers efficiency was calculated by serial sample dilution. Targeted gene expressions were normalized by Cyclophilin reference gene.

Cyclophilin-Fwd: AAGAAGATCACCATTTCCGACT,
Cyclophilin-Rev: TTA CAG GAC ATT GCG AGC,
MyH3-Fwd: AGAGGAGAAGGCCAAAAGG,
MyH3-Rev CCTTCCAGCTCAAACTCCAG,
Myogenin-Fwd: GAAAGTGAATGAGGCCTTCG,
Myogenin-Rev: ACGATGGACGTAAGGGAGTG,
Pax7-Fwd: CTGGATGAGGGCTCAGATGT,
Pax7-Rev: GGTTAGCTCCTGCCTGCTTA,
Grem1-Fwd: AAGCGAGATTGGTGCAAAACT,
Grem1-Rev: GAAGCGGTTGATGATAGTGCG,
Prkaa2-Fwd: CAGGCCATAAAGTGGCAGTTA
Prkaa2-Rev:AAAAGTCTGTCGGAGTGCTGA,
Prkag3-Fwd: ATCTCTCCCAATGACAGCCTG,
Prkag3-Rev: TAGCCGCTTGTGTGTGAGTAT,
Prkaa1-Fwd TACTCAACCGGCAGAAGATTCG,
Prkaa1-Rev AGACGGCGGCTTTCCTTTT;
Bmp4-Fwd: TAGCAAGAGCGCCGTCATTC,
Bmp4-Rev: CTGGTCCCTGGGATGTTCTC,
Id1-Fwd: TCCTGCAGCATGTAATCGAC,
Id1-Rev: GATCGTCGGCTGGAACAC,
Tbp-Fwd: CCCCACAACTCTTCCATTCT,
Tbp-Rev: GCAGGAGTGATAGGGGTCAT.

### Transcriptomic analysis

Total RNA from primary myoblasts were isolated using TRIzol Reagent from Life Technologies according to the manufacturer’s protocol. The purity and the quality of the RNA were performed by Bioanalyzer 2100, using Agilent RNA6000 nano chip kit (Agilent Technologies) at the Cochin Institute Genom’IC facility. Total RNA was reverse transcribed using the Ovation PicoSL WTA System (Nugen). The products cDNA products then hybridized to GeneChip Mouse Gene 2.1 ST (Affymetrix). The Affymetrix data were normalized using R software^79^ and gene expression levels were compared using one-way ANOVA. Differentially expressed genes were selected with *p*<0.05 significance. Functional analysis of the gene expression levels was interrogated with Ingenuity Pathway (Ingenuity Systems, http://www.ingenuity.com).

### Western-blot analysis

Cells were lysed with RIPA buffer (Sigma-Aldrich) supplemented with Protease and Phosphatase Inhibitor (Thermo Fisher). Protein amounts were quantified by BCA Protein Assay Kit (Pearce). In each condition, 30μg of proteins was prepared with home-made Laemmli. Samples were ran on NuPAGE 4-12% BIS-TRIS gels (Life Technologies) and transferred on nitrocellulose membranes. After blocking in 5% milk and 0.1% Tween-20 in TBS, membranes were incubated with primary antibodies overnight at 4°C. Secondary antibodies were then incubated on the membrane for 1 hr at room temperature. The following antibodies were used: Gremlin C-terminal (Abcam), phospho-Ampkα1/2 (Thr172, #2535), Ampkα1/2 (#2532), phospho-Akt (Ser473, #4060), Akt (#4691) or β-Actin (D6A8, #8457) (all fivefrom Cell Signaling Technology) antibodies. Signals were detected using SuperSignalWest Pico Chemiluminescent Substrate (Thermo Scientific) with ImageQuant TL LAS 4000 from (GE Healthcare) or Bio-Rad ChemiDoc with densitometry analysis performed using Imagelab software version 3.0 (Bio-Rad). The primary antibodies used were as follows: SMAD1 (Cell Signaling), phosphorylated SMAD1 (Cell Signaling) and GAPDH (Sigma).

### Image acquisition and quantitative analysis

Immunofluorescent staining was analyzed with an Olympus BX63F microscope, AZ100 Nikon Macroscope and EVOS FL Cell Imaging System. Fluorochromes used were Alexa Fluor 488 and Alexa Fluor 546 from Life Technologies. NIS-Element (Nikon), Metamorph (Molecular Devices), and Photoshop software (Adobe) were used for image acquisition. Quantifications were performed with ImageJ software.

### Electrical Pulse Stimulation (EPS)

Differentiated C2C12 muscle cells were subjected to EPS at 30v, 2 ms, 1 Hz for 30 min twice with overnight resting between first and second stimulation. Cells were harvested immediately, 1, 3, 4, 5 and 6 hrs after last stimulation. Control and non-stimulated cells were exposed to electrodes without electrical stimulation. RNA and DNA were harvested by AllPrep DNA/RNA/miRNA universal kit. cDNA was synthesized from RNA using iScript cDNA synthesis kit. Then, mRNA expression was quantified by qPCR and normalized by cDNA concentration.

To capture methylated DNA, genomic DNA was sheared with Bioruptor Plus sonication and then methylated DNA was captured using

MethylCollector ultra kit (Active Motif). Captured and non-captured DNA were purified by MinElute PCR purification kit. Methylation levels of the Grem1-DMR were then quantified by qPCR using primers spanning DMR region. Data were normalized by non-methylated DNA level.

*Grem1-DMR* fwd primer: GCCTCATGAGCTTTAAGTAGGGA;

*Grem1-DMR* rev primer: CTGCAACAAAGTGAAAAGACAATCT.

### Generation of the C2C12 MS2-CAS cell line

Lentiviruses containing MS2-P65-HSF1_Hygro or dCAS-VP64_Blast shRNA were produced in HEK293 by also transfecting cells with pMD2.G (viral envelope) and psPAX2 (packaging) at the *ratio* 0.5 μg pMD2.G/1.5 μg psPAX2/2 μg shRNA. All plasmids were purchased from Addgene. Transfection was realized with Superfect (Qiagen). Lentivirus-containing medium was collected 48 hrs after transfection and stored at −80°C. C2C12 MS2-CAS cell line was generated by sequential transduction of the mouse myogenic C2C12 cells with both of the lentiviruses and selection using hygromycin 0.2 mg/ml and blasticidin 4 μg/ml (Sigma-Aldrich) in growth medium. Integration of plasmids in the genome of newly generated C2C12 MS2-CAS cells was controlled using the following primers:

MS2-p65-HSF1 Fwd 5’-GGGCTCCTCAAAGACGGTAA-3’;
MS2-p65-HSF1 Rev 5’-ATACCTGAGTTAGCGGCGAT-3’;
dCas9-VP64 Fwd 5’-ACGCTAATCTGGACAAAGTGC-3’;
dCas9-VP64 Rev 5’-CCTGCTCTCTGATGGGCTTAT-3’;.

Viability of transfected cells was assessed using a NucleoCounter NC-200.

### Over-expression of *Grem1* using C2C12 MS2-CAS cells

CRISPR/Cas9 Synergistic Activation Mediator (SAM) is an engineered protein complex for the transcriptional activation of endogenous genes^47^. Mouse *Grem1* sgRNA sequence was obtained using the Cas9 SAM sgRNA design tool available online (http://sam.genome-engineering.org/database/). Potential off-targets were identified by a CRISPR analysis tool (http://crispr.mit.edu/) and three of them were considered as presenting a high risk to interfere with *Grem1* over-expression because located in a gene promoter. Grem1 sgRNA was incorporated into the sgRNA(MS2) cloning backbone (Addgene #61424) following the protocol available online *via* the Zhang lab website (http://sam.genome-engineering.org/protocols/). Correct insertion of the sgRNA sequence in the cloning backbone was checked by Sanger DNA sequencing (GENEWIZ). C2C12 MS2-CAS cells were grown in medium composed by high-glucose DMEM (Invitrogen) supplemented with FBS (Sigma-Aldrich) 10% (V/V) and Pen Strep 1% (Invitrogen), and maintained in 5% CO_2_ at 37°C. When 80% confluent, cells were transfected using *Grem1* sgRNA-containing plasmid or the cloning backbone (empty vector) using the TransIT-X2 Dynamic Delivery System (Mirus). After 24 hrs, the transfection medium was removed and replaced by differentiation medium. Viability of transfected cells was assessed in proliferative myoblasts using a NucleoCounter NC-200.

### Proteomics analysis

Cells were quickly washed twice with cold 1x PBS. Lysis buffer (6 M guanidium hydrochloride and 25 mM Tris (pH 8) containing 5 mM tris (2-carboxyethyl) phosphine (TCEP) and 5.5 mM chloroacetamide (CAA)) was added to culture plates. Cells were scraped and samples were boiled at 99 °C for 10 min followed by sonication using a tip. Proteins were enzymatically digested using Lys-C (Wako Chemicals, Richmond, VA, UsA) at a 1:100 (w/w) ratio for 3 h at 37 °C. Samples were further diluted with 50 mM ammonium bicarbonate to 2M guanidinium concentration, and trypsin was added at 1:100 (w/w) to generate tryptic peptides overnight at 37 °C. After digestion, samples were acidified with trifluoroacetic acid (TFA, 1% final) and centrifuged to remove debris. Peptides were desalted on a C18 StageTips^86^. Peptides were measured using LC-MS instrumentation consisting of an Easy nanoflow HPLC system (Thermo Fisher Scientific, Bremen, Germany) coupled via a nanoelectrospray ion source (Thermo Fischer Scientific, Bremen, Germany) to a Q Exactive HF mass spectrometer^87^. Purified peptides were separated on 50 cm C10 column (inner diameter 75 μm, 1.8 μm beads, Dr. Maisch GmbH, Germany). Peptides were loaded onto the column with buffer A (0.5% formic acid) and eluted with a 120 min linear gradient from 5-40% buffer B (80% acetonitrile, 0.5% formic acid). After the gradient, the column was washed with 80% buffer B and reequilibrated with buffer A. Mass spectra were acquired in a data-dependent manner with automatic switching between MS and MS/MS using a top 12 method. MS spectra were acquired in the Orbitrap analyzer with a mass range of 300-1750 m/z and 60,000 resolutions at m/z 200. HCD peptide fragment acquired at 28 normalized collision energy were analyzed at high resolution in the Orbitrap analyzer.

Raw MS files were analyzed using MaxQuant version 1.5.0.38^88^ (http://www.maxquant.org). MS/MS spectra were searched by the Andromeda search engine (integrated into MaxQuant) against the decoy UniProt-human database with forward and reverse sequences. In the main Andromeda search precursor, mass and fragment mass were matched with an initial mass tolerance of 6 ppm and 20 ppm, respectively. The search included variable modifications of methionine oxidation and N-terminal acetylation and fixed modification of carbamidomethyl cysteine. Minimal peptide length was set to 7 amino acids, and a maximum of two miscleavages was allowed. The FDR was 0.01 for peptide and protein identifications. When all identified peptides were shared between two proteins, results were combined and reported as one protein group. Matches to the reverse database were excluded. Protein quantification was based on the MaxLFQ algorithm integrated into the MaxQuant software^89^.

### Palmitate oxidation assays

Fully differentiated C2C12 MS2-CAS myotubes were starved in serum-free DMEM for 2 hrs before a 3-hr incubation in glucose-free or low-glucose DMEM/fatty acid-free BSA (Sigma-Aldrich) 0.2% supplemented with a mixture of cold and radiolabeled palmitate (Palmitic Acid, [9,10-^3^H(N)]; 0.5 μCi/ml; PerkinElmer) at final concentration of 100 or 200 μM^14,15^. Incubation media were collected and 200 μl were mixed with 800 μl Tris-HCl [pH 7.5] 0.02M/activated charcoal slurry (Sigma-Aldrich) 10% for 1 hr on a rotational shaker^90^. After a 15-min centrifugation at full speed at room temperature, 300 μl of each supernatant were mixed with 4 ml of Ultima Gold liquid scintillation cocktail (PerkinElmer) and run on a Hidex SL300 counter, with 600 sec counting. Myotubes were harvested in lysis buffer for protein purification and DPM values were normalized by total protein content.

For palmitate oxidation assays in cells from wild type or AMPKα1-muscle knockout (mKO) mice, muscle progenitor cells were isolated from hindlimb muscles of 4-week old wild type or AMPKα1-muscle knockout (mKO) mice as previously described^91^. Mouse muscle progenitor cells were cultured in DMEM/F12 supplemented with 1% Penicillin and Streptomycin (P/S), 20% fetal bovine serum (FBS, Sigma Aldrich) and 2% Ultroser G (Pall Life Sciences) at 37°C in a humified atmosphere containing 5% CO_2_. Myogenic differentiation was initiated 6 hrs after seeding cells onto Matrigel (BD Life Sciences) coated cell culture dishes (30,000 cells/cm^2^) and switching to myogenic differentiation media (DMEM/F12, 1% Pen/Strep, 2% horse serum). Three days after induction of differentiation, cells were incubated for 3 hrs in glucose-free DMEM/fatty acid-free BSA (Sigma-Aldrich) 0.2%, supplemented with a mixture of cold and radiolabeled palmitate (Palmitic Acid, [9,10-3H(N)]; 0.5 μCi/ml; PerkinElmer) at a final concentration of 600 μM. Vehicle (4 mM HCl-supplemented 0.1 % bovine serum albumin (Sigma-Aldrich)), 2 mM 5-aminoimidazole-4-carboxamide ribonucleotide (AICAR, Toronto Research Chemicals), or 500 ng/mL mouse recombinant Gremlin 1 (R&D Biosciences) was added to the assay media, and palmitate oxidation was measured as described above.

### Glucose uptake assay

Fully differentiated C2C12 MS2-CAS myotubes were starved in serum-and glucose-free DMEM supplemented with BSA 0.1% for 2.5 hrs before a 10-min incubation in Krebs-Ringer Phosphate HEPES (KRPH) buffer [pH 7.4]/BSA 0.1% supplemented with a mixture of cold deoxyglucose (2-Deoxy-D-glucose; Sigma-Aldrich) and radiolabeled deoxyglucose (Deoxy-D-glucose, 2[1,2-^3^H(N)]; 0.5 μCi/ml; PerkinElmer) for a final concentration of 100 μM. Then, myotubes were harvested in lysis buffer for protein purification and a 150-μl fraction of protein extract was mixed with 4 ml of Ultima Gold liquid scintillation cocktail and run on a Hidex SL300 counter, with 300 sec counting. DPM values were normalized by protein content.

### Glucose oxidation assay

Fully differentiated C2C12 MS2-CAS myotubes were starved in serum-and glucose-free DMEM for 2 hrs before a 3-hr incubation in low-glucose DMEM/fatty acid-free BSA 0.2% supplemented with radiolabeled glucose (Glucose, D-[3-^3^H]; 1 μCi/ml; PerkinElmer). Then, cell media and lysates were collected and processed as described above for palmitate oxidation assays.

### Glycogen synthesis assay

Fully differentiated C2C12 MS2-CAS myotubes were starved in serum-and glucose-free DMEM for 2 hrs before a 3-hr incubation in low-glucose DMEM/fatty acid-free BSA 0.2%. Then, cells were harvested in 150 μl HCl 1M and heated at 95°C for 2 hrs. After heating, HCl was immediately neutralized in 75 μl NaOH 2M. Samples were centrifuged at full speed at 4°C and 20 μl of the supernatants were mixed with 250 μl of a solution composed by Tris-HCL [pH 7.4], MgCl_2_ 500 mM, ATP disodium salt 5 nM, NADP 3 nM (all from Sigma-Aldrich) and Hexokinase + Glucose-6-Phosphate dehydrogenase at dilution 1:200 (Roche), for 15 min at 25°C. Absorbance was read at 340 nm and concentrations were normalized by protein content.

### Oil Red O staining

Three days after induction of adipogenic differentiation, cells were fixed with 4% formaldehyde for 15 min at room temperature; then neutral lipids were stained with Oil Red O (Sigma-Aldrich) 0.5% in isopropanol/distilled water (3V/2V) for 60 min at room temperature. Stained cells were visualized using a Primovert microscope (Carl Zeiss Microscopy). Cells were then rinsed with water to remove unbound dye and incubated 10 min with ethanol to elute the Oil Red O. Intracellular lipid accumulation was quantified by spectrophotometric analysis of the eluates at 540 nm.

### Statistics

Statistical analyses were performed using GraphPad Prism software version s6 and 7 (GraphPad Software Inc). For the experiments done on human primary muscle cells, we conducted a Student’s t-test for paired samples to compare pre-and post-training or control and EPS samples. For the other *in vitro* experiments, we conducted a Student’s t-test for paired samples or a non-parametric Wilcoxon signed-rank test when normality failed (*p*>0.05) or a two-way repeated measures ANOVA with Sidak’s multiple comparisons test to compare the effect of two variables (time x *Grem1* or *Grem1* x treatment) on *Grem1* sgRNA-and empty vector-transfected cells. Ordinary one-way ANOVA with Tukey’s multiple comparison test were used to compare multiple treatments (Grem1, Bmp4, Grem1+Bmp4). A two-tailed non-parametric Spearman test was conducted for correlation analyses. For in vivo experiments, we conducted Student’s t-test for paired samples. A two-tailed non-parametric Spearman test was conducted for correlation analyses. Statistical analyses of differences on mRNA expression were performed on log-transformed values. We considered that the difference between two conditions was statistically significant when *p≤0.05, **p≤0.01 or ***p≤0.001 and reported a tendency when 0.05<p<0.1. Data are expressed as mean ± SD or ±SEM as indicated in figure legends.

## End Matter

### Author Contributions and Notes

O.F., L.G., A.P., P.P., D.A., C.B., I.C., A.T., A.B., E.A., P.M., M.R., C.L., K.C. and F.L.G. contributed to data acquisition. O.F., L.G., A.P., P.P., D.A., C.B., I.C., A.T., A.B., E.A., P.M., M.R., A.L., A.D., C.M., V.B., R.M., F.L.G. and R.B. contributed to data analysis. O.F., V.B., C.M., R.M., F.L.G. and R.B. contributed to the study design. All authors contributed to data interpretation and manuscript drafting and approved the final version of the manuscript. F.L.G. and R.B. are the guarantors of this work and, as such, had full access to all the data in the study and takes responsibility for the integrity of the data and the accuracy of the data analysis.

The authors declare no conflict of interest.

### Funding Sourcess

This work was supported by funding from the INSERM, the CNRS, the Agence Nationale pour la Recherche (ANR-14-CE11-0026, ANR-12-JSV2-0003, ANR-17-CE12-0010-02), and the Association Française contre les Myopathies/AFM Telethon. A.P. was supported by a PhD fellowship from the Ministère de la Recherche et de l’Enseignement and a fellowship from the Fondation ARC pour la Recherche sur le Cancer.

O.F. was recipient of a research grant from the Danish Diabetes Academy supported by the Novo Nordisk Foundation.

## Acknowledgments

We would like to acknowledge Morten Lundh and Louise Larsson, Novo Nordisk Foundation Center for Basic Metabolic Research, University of Copenhagen, for assistance with the generation of the C2C12 MS2-CAS cell line and with palmitate oxidation assays, respectively, as well as The Danish National High-Throughput DNA Sequencing Centre, University of Copenhagen, for sequencing the samples and the Core Facility for Integrated Microscopy, University of Copenhagen, for assistance with immunofluorescence experiments. We thank Rebeca Soria Romero for assistance with sample preparation for proteomics analysis. The authors also acknowledge B. Durel and P. Bourdoncle of the Cochin Imaging Facility. We thank members of the Cochin Institute Genom’IC Facility (S. Jacques and F. Dumont) and members of the UMS37 CyPS Facility (C. Blanc and B. Hoareau-Coudert) for technical support. The Novo Nordisk Foundation Center for Basic Metabolic Research is an independent research center at the University of Copenhagen partially funded by an unrestricted donation from the Novo Nordisk Foundation.

---

**Supplementary Fig. 1.**
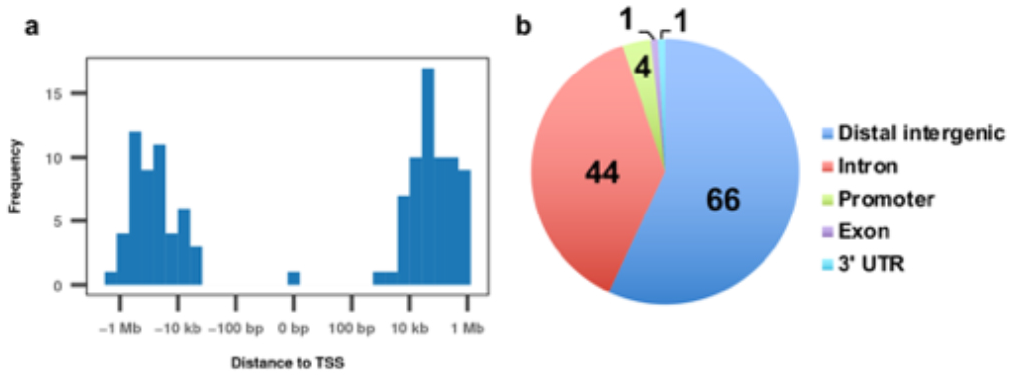
Genome-wide DNA methylation results. Distribution of identified DMRs according to their distance to nearest transcription start site (TSS) and b, amongst different gene features.

**Supplementary Fig. 2.**
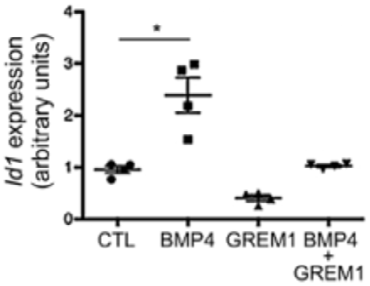
GREM1 prevents the increase in mRNA expression of the BMP target genes Id1. RT-qPCR analysis of the *Bmp4* target gene *Id1* expression in primary myocytes stimulated with recombinant BMP4, GREM1 or both. n = 4. Error bars indicate SD. \**p*<0.05; \*\**p*<0.01; \*\*\**p*< 0.001.

**Supplementary Fig. 3.**
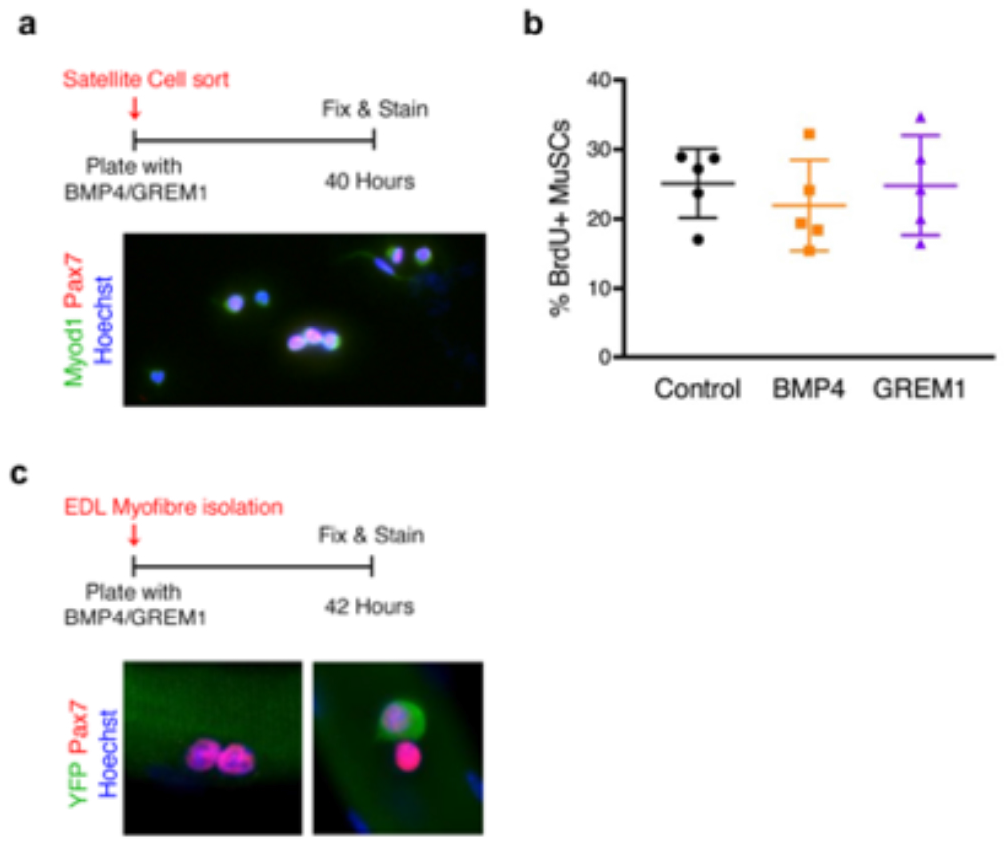
GREM1 stimulates asymmetric commitment of satellite stem cells. **a**, Schematic of experimental setup. Satellite cells were sorted and plated *in vitro* with growth medium containing recombinant proteins, fixed after 40 hrs and stained for Pax7 (red) and Myod1 (green). **b**, BrdU incorporation assay. n = 5. **c**, Schematic of experimental setup. Single myofibres from Myf5-Cre*ROSA-YFP mice were separated and cultured in floating conditions for 42 hrs in growth medium containing recombinant proteins. Myofibers were fixed and stained for Pax7 (red) and YFP (green).

**Supplementary Fig. 4.**
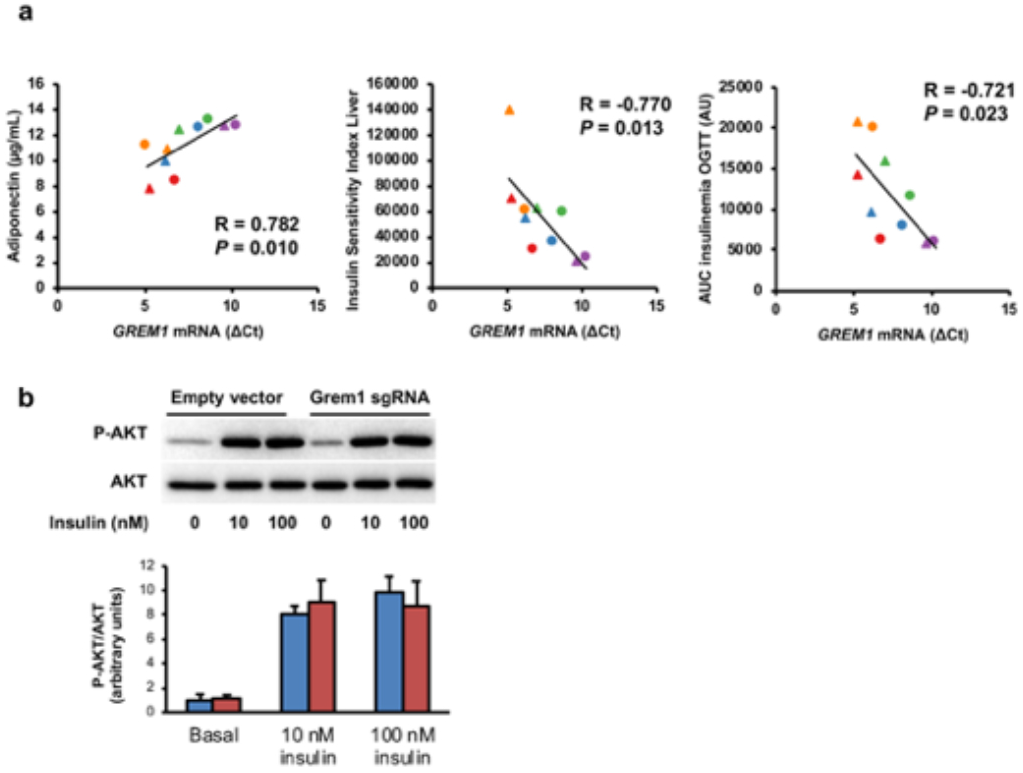
Characterization of endogenous Grem1-overexpressing cells. Before reaching confluence, C2C12 MS2-CAS cells were transfected with Grem1 sgRNA or empty vector for 24 hrs and switched to myogenic differentiation medium. **a**, Control for possible gRNA off-target effects by RT-qPCR. **b**, Representative photomicrographs were taken and cell viability quantified. **c**, Cells were harvested every day from confluence and analysis of *Grem1* mRNA expression was made by RT-qPCR. Each individual dot in the graph represents the mean of 2^-ΔCt^ duplicate values from an independent experiment. Vertical axis scale is logarithmic base 10. ^§^p<0.001. **d**, Cell media was collected at day 3 of differentiation for protein extraction and analysis of GREM1 expression by western-blot. n = 3. \*\**p*<0.005.

**Supplementary Fig. 5.**
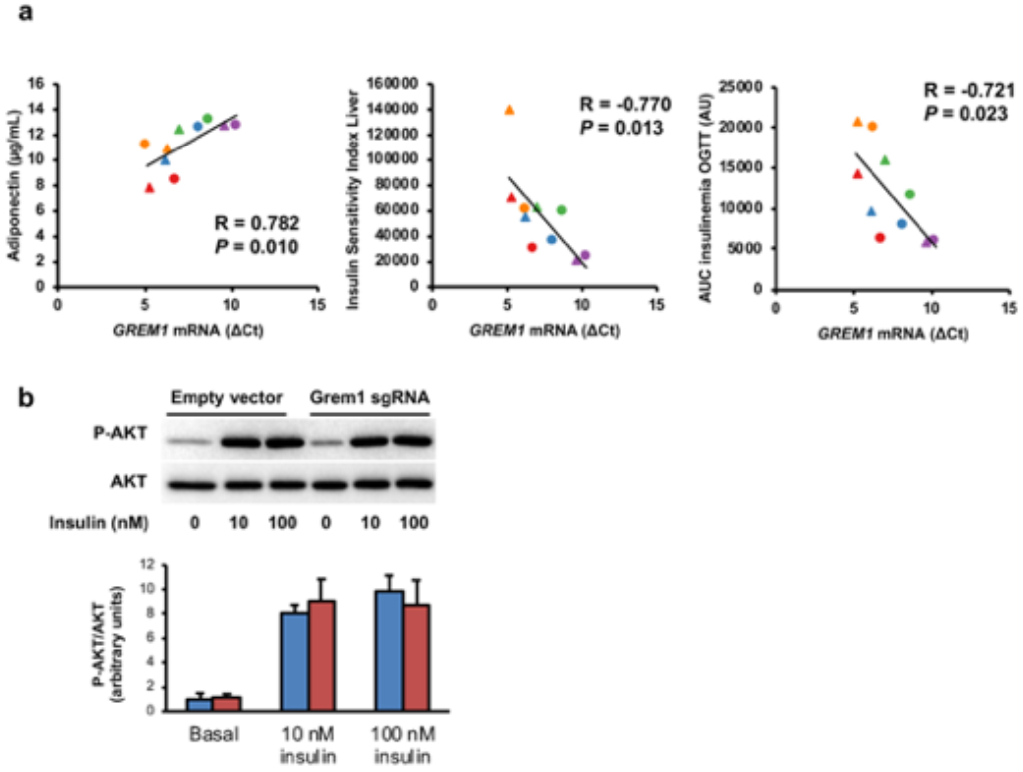
Correlation analysis between GREM1 expression and clinical parameters of obese subjects. **a**, Correlation analyses were performed between ΔCt values, determined by RT-qPCR, and clinical and biochemical variables collected from the obese subjects pre-and post-training (n = 10) described in (Bourlier et al., 2013). Circle = Pretraining; triangle = Post-training. **b**, Myotubes were incubated for 10 min in presence or not of insulin 10 nM or 100 nM for the analysis of P-AKT and AKT expression by Western-blot. Duplicate protein extracts from each experiment were equally mixed according to their molarity before loading the gel. n = 4. The graph represents mean values. Error bars are SEM.

**Supplementary Fig. 6.**
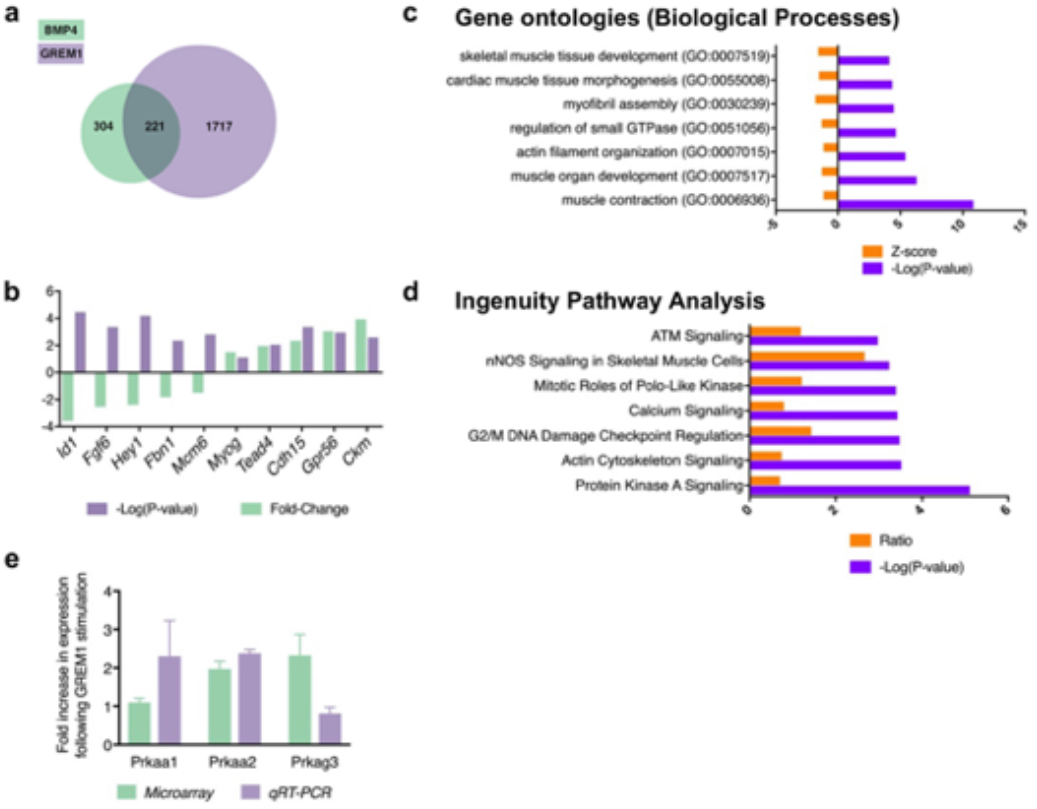
Transcriptomic analysis of muscle cells exposed to GREM1 highlights the AMPK pathway. **a**, Differentially expressed genes after overexpression of BMP4 (green) or GREM1 (purple) in serum-starved mouse myoblasts. **b**, Difference in expression of ten selected transcripts between GREM1-stimulated cells compared with BMP4-stimulated cells. GREM1 increases the expression of genes related to muscle cell differentiation, while BMP4 stimulates the expression of genes involved in proliferation. **c**, Gene Ontology analysis of biological processes over-represented in GREM1-stimulated cells compared with mock-treated cells (top 7 terms). **d**, Ingenuity Pathway Analysis of differentially expression genes in GREM1-stimulated cells compared with mock-treated cells. **e**, Gene expression analysis of AMPK-related genes following GREM1 stimulation. n = 3 or more for any experimental point.

## References

1 Pedersen, B. K. & Febbraio, M. A. Muscles, exercise and obesity: skeletal muscle as a secretory organ. Nat Rev Endocrinol 8, 457–465, doi:10.1038/nrendo.2012.49 (2012).

2 DeFronzo, R. A. Lilly lecture 1987. The triumvirate: beta-cell, muscle, liver. A collusion responsible for NIDDM. Diabetes 37, 667–687, doi:10.2337/diab.37.6.667 (1988).

3 Chazaud, B. Inflammation during skeletal muscle regeneration and tissue remodeling: application to exercise-induced muscle damage management. Immunol Cell Biol 94, 140–145, doi:10.1038/icb.2015.97 (2016).

4 Relaix, F. & Marcelle, C. Muscle stem cells. Curr Opin Cell Biol 21, 748–753, doi:10.1016/j.ceb.2009.10.002 (2009).

5 Appell, H. J., Forsberg, S. & Hollmann, W. Satellite cell activation in human skeletal muscle after training: evidence for muscle fiber neoformation. Int J Sports Med 9, 297–299, doi:10.1055/s-2007-1025026 (1988).

6 Charifi, N., Kadi, F., Feasson, L. & Denis, C. Effects of endurance training on satellite cell frequency in skeletal muscle of old men. Muscle Nerve 28, 87–92, doi:10.1002/mus.10394 (2003).

7 Petrella, J. K., Kim, J. S., Mayhew, D. L., Cross, J. M. & Bamman, M. M. Potent myofiber hypertrophy during resistance training in humans is associated with satellite cell-mediated myonuclear addition: a cluster analysis. J Appl Physiol (1985) 104, 1736–1742, doi:10.1152/japplphysiol.01215.2007 (2008).

8 Bouzakri, K. et al. Reduced activation of phosphatidylinositol-3 kinase and increased serine 636 phosphorylation of insulin receptor substrate-1 in primary culture of skeletal muscle cells from patients with type 2 diabetes. Diabetes 52, 1319–1325, doi:10.2337/diabetes.52.6.1319 (2003).

9 Bouzakri, K. & Zierath, J. R. MAP4K4 gene silencing in human skeletal muscle prevents tumor necrosis factor-alpha-induced insulin resistance. J Biol Chem 282, 7783–7789, doi:10.1074/jbc.M608602200 (2007).

10 Gaster, M., Petersen, I., Hojlund, K., Poulsen, P. & Beck-Nielsen, H. The diabetic phenotype is conserved in myotubes established from diabetic subjects: evidence for primary defects in glucose transport and glycogen synthase activity. Diabetes 51, 921–927, doi:10.2337/diabetes.51.4.921 (2002).

11 Henry, R. R., Abrams, L., Nikoulina, S. & Ciaraldi, T. P. Insulin action and glucose metabolism in nondiabetic control and NIDDM subjects. Comparison using human skeletal muscle cell cultures. Diabetes 44, 936–946, doi:10.2337/diab.44.8.936 (1995).

12 Henry, R. R. et al. Glycogen synthase activity is reduced in cultured skeletal muscle cells of non-insulin-dependent diabetes mellitus subjects. Biochemical and molecular mechanisms. J Clin Invest 98, 1231–1236, doi:10.1172/JCI118906 (1996).

13 Gaster, M., Rustan, A. C., Aas, V. & Beck-Nielsen, H. Reduced lipid oxidation in skeletal muscle from type 2 diabetic subjects may be of genetic origin: evidence from cultured myotubes. Diabetes 53, 542–548, doi:10.2337/diabetes.53.3.542 (2004).

14 Ukropcova, B. et al. Dynamic changes in fat oxidation in human primary myocytes mirror metabolic characteristics of the donor. J Clin Invest 115, 1934–1941, doi:10.1172/JCI24332 (2005).

15 Bourlier, V. et al. Enhanced glucose metabolism is preserved in cultured primary myotubes from obese donors in response to exercise training. J Clin Endocrinol Metab 98, 3739–3747, doi:10.1210/jc.2013-1727 (2013).

16 Costford, S. R. et al. Gain-of-function R225W mutation in human AMPKgamma(3) causing increased glycogen and decreased triglyceride in skeletal muscle. PLoS One 2, e903, doi:10.1371/journal.pone.0000903 (2007).

17 Crawford, S. A. et al. Naturally occurring R225W mutation of the gene encoding AMP-activated protein kinase (AMPK)gamma(3) results in increased oxidative capacity and glucose uptake in human primary myotubes. Diabetologia 53, 1986–1997, doi:10.1007/s00125-010-1788-7 (2010).

18 Bharathy, N. & Taneja, R. Methylation muscles into transcription factor silencing. Transcription 3, 215–220, doi:10.4161/trns.20914 (2012).

19 Sousa-Victor, P., Munoz-Canoves, P. & Perdiguero, E. Regulation of skeletal muscle stem cells through epigenetic mechanisms. Toxicol Mech Methods 21, 334–342, doi:10.3109/15376516.2011.557873 (2011).

20 Cedar, H. & Bergman, Y. Linking DNA methylation and histone modification: patterns and paradigms. Nat Rev Genet 10, 295–304, doi:10.1038/nrg2540 (2009).

21 Barres, R. et al. Acute exercise remodels promoter methylation in human skeletal muscle. Cell Metab 15, 405–411, doi:10.1016/j.cmet.2012.01.001 (2012).

22 Fabre, O. et al. Exercise training alters the genomic response to acute exercise in human adipose tissue. Epigenomics 10, 1033–1050, doi:10.2217/epi-2018-0039 (2018).

23 Denham, J., Marques, F. Z., O’Brien, B. J. & Charchar, F. J. Exercise: putting action into our epigenome. Sports Med 44, 189–209, doi:10.1007/s40279-013-0114-1 (2014).

24 Horsburgh, S., Robson-Ansley, P., Adams, R. & Smith, C. Exercise and inflammation-related epigenetic modifications: focus on DNA methylation. Exerc Immunol Rev 21, 26–41 (2015).

25 Nitert, M. D. et al. Impact of an exercise intervention on DNA methylation in skeletal muscle from first-degree relatives of patients with type 2 diabetes. Diabetes 61, 3322–3332, doi:10.2337/db11-1653 (2012).

26 Lindholm, M. E. et al. An integrative analysis reveals coordinated reprogramming of the epigenome and the transcriptome in human skeletal muscle after training. Epigenetics 9, 1557–1569, doi:10.4161/15592294.2014.982445 (2014).

27 Eden, E., Navon, R., Steinfeld, I., Lipson, D. & Yakhini, Z. GOrilla: a tool for discovery and visualization of enriched GO terms in ranked gene lists. BMC Bioinformatics 10, 48, doi:10.1186/1471-2105-10-48 (2009).

28 McLean, C. Y. et al. GREAT improves functional interpretation of cis-regulatory regions. Nat Biotechnol 28, 495–501, doi:10.1038/nbt.1630 (2010).

29 Pavel, E., Zhao, W., Powell, K. A., Weinstein, M. & Kirschner, L. S. Analysis of a new allele of limb deformity (ld) reveals tissue-and age-specific transcriptional effects of the Ld Global Control Region. Int J Dev Biol 51, 273–281, doi:10.1387/ijdb.062249ep (2007).

30 Zuniga, A. et al. Mouse limb deformity mutations disrupt a global control region within the large regulatory landscape required for Gremlin expression. Genes Dev 18, 1553–1564, doi:10.1101/gad.299904 (2004).

31 Liu, Y. et al. Six1 regulates MyoD expression in adult muscle progenitor cells. PLoS One 8, e67762, doi:10.1371/journal.pone.0067762 (2013).

32 Le Grand, F. et al. Six1 regulates stem cell repair potential and selfrenewal during skeletal muscle regeneration. J Cell Biol 198, 815–832, doi:10.1083/jcb.201201050 (2012).

33 Hsu, D. R., Economides, A. N., Wang, X., Eimon, P. M. & Harland, R. M. The Xenopus dorsalizing factor Gremlin identifies a novel family of secreted proteins that antagonize BMP activities. Mol Cell 1, 673–683 (1998).

34 Merino, R. et al. The BMP antagonist Gremlin regulates outgrowth, chondrogenesis and programmed cell death in the developing limb. Development 126, 5515–5522 (1999).

35 Topol, L. Z., Modi, W. S., Koochekpour, S. & Blair, D. G. DRM/GREMLIN (CKTSF1B1) maps to human chromosome 15 and is highly expressed in adult and fetal brain. Cytogenet Cell Genet 89, 79–84, doi:10.1159/000015568 (2000).

36 Frank, N. Y. et al. Regulation of myogenic progenitor proliferation in human fetal skeletal muscle by BMP4 and its antagonist Gremlin. J Cell Biol 175, 99–110, doi:10.1083/jcb.200511036 (2006).

37 Walsh, D. W., Godson, C., Brazil, D. P. & Martin, F. Extracellular BMP-antagonist regulation in development and disease: tied up in knots. Trends Cell Biol 20, 244–256, doi:10.1016/j.tcb.2010.01.008 (2010).

38 Katagiri, T. et al. Bone morphogenetic protein-2 inhibits terminal differentiation of myogenic cells by suppressing the transcriptional activity of MyoD and myogenin. Exp Cell Res 230, 342–351, doi:10.1006/excr.1996.3432 (1997).

39 Murray, S. S., Murray, E. J., Glackin, C. A. & Urist, M. R. Bone morphogenetic protein inhibits differentiation and affects expression of helix-loop-helix regulatory molecules in myoblastic cells. J Cell Biochem 53, 51–60, doi:10.1002/jcb.240530107 (1993).

40 Korchynskyi, O. & ten Dijke, P. Identification and functional characterization of distinct critically important bone morphogenetic protein-specific response elements in the Id1 promoter. J Biol Chem 277, 4883–4891, doi:10.1074/jbc.M111023200 (2002).

41 Kuang, S., Kuroda, K., Le Grand, F. & Rudnicki, M. A. Asymmetric self-renewal and commitment of satellite stem cells in muscle. Cell 129, 999–1010, doi:10.1016/j.cell.2007.03.044 (2007).

42 Bentzinger, C. F., Wang, Y. X., Dumont, N. A. & Rudnicki, M. A. Cellular dynamics in the muscle satellite cell niche. EMBO Rep 14, 1062–1072, doi:10.1038/embor.2013.182 (2013).

43 Le Grand, F., Jones, A. E., Seale, V., Scime, A. & Rudnicki, M. A. Wnt7a activates the planar cell polarity pathway to drive the symmetric expansion of satellite stem cells. Cell Stem Cell 4, 535–547, doi:10.1016/j.stem.2009.03.013 (2009).

44 Wang, J. et al. Bmp signaling regulates myocardial differentiation from cardiac progenitors through a MicroRNA-mediated mechanism. Dev Cell 19, 903–912, doi:10.1016/j.devcel.2010.10.022 (2010).

45 Abou-Khalil, R., Le Grand, F. & Chazaud, B. Human and murine skeletal muscle reserve cells. Methods Mol Biol 1035, 165–177, doi:10.1007/978-1-62703-508-8_14 (2013).

46 Gustafson, B. et al. BMP4 and BMP Antagonists Regulate Human White and Beige Adipogenesis. Diabetes 64, 1670–1681, doi:10.2337/db14-1127 (2015).

47 Konermann, S. et al. Genome-scale transcriptional activation by an engineered CRISPR-Cas9 complex. Nature 517, 583–588, doi:10.1038/nature14136 (2015).

48 Coffey, V. G. & Hawley, J. A. The molecular bases of training adaptation. Sports Med 37, 737–763, doi:10.2165/00007256-200737090-00001 (2007).

49 Kusuhara, K., Madsen, K., Jensen, L., Hellsten, Y. & Pilegaard, H. Calcium signalling in the regulation of PGC-1alpha, PDK4 and HKII mRNA expression. Biol Chem 388, 481–488, doi:10.1515/BC.2007.052 (2007).

50 Ojuka, E. O., Jones, T. E., Han, D. H., Chen, M. & Holloszy, J. O. Raising Ca2+ in L6 myotubes mimics effects of exercise on mitochondrial biogenesis in muscle. FASEB J 17, 675–681, doi:10.1096/fj.02-0951com (2003).

51 Ojuka, E. O. et al. Intermittent increases in cytosolic Ca2+ stimulate mitochondrial biogenesis in muscle cells. Am J Physiol Endocrinol Metab 283, E1040–1045, doi:10.1152/ajpendo.00242.2002 (2002).

52 Louche, K. et al. Endurance exercise training up-regulates lipolytic proteins and reduces triglyceride content in skeletal muscle of obese subjects. J Clin Endocrinol Metab 98, 4863–4871, doi:10.1210/jc.2013-2058 (2013).

53 Theret, M. et al. AMPKalpha1-LDH pathway regulates muscle stem cell self-renewal by controlling metabolic homeostasis. EMBO J 36, 1946–1962, doi:10.15252/embj.201695273 (2017).

54 Jacques, M. et al. Epigenetic changes in healthy human skeletal muscle following exercise-a systematic review. Epigenetics 14, 633–648, doi:10.1080/15592294.2019.1614416 (2019).

55 Seaborne, R. A. et al. Human Skeletal Muscle Possesses an Epigenetic Memory of Hypertrophy. Sci Rep 8, 1898, doi:10.1038/s41598-018-20287-3 (2018).

56 Robinson, M. M. et al. Enhanced Protein Translation Underlies Improved Metabolic and Physical Adaptations to Different Exercise Training Modes in Young and Old Humans. Cell Metab 25, 581–592, doi:10.1016/j.cmet.2017.02.009 (2017).

57 Pattamaprapanont, P., Garde, C., Fabre, O. & Barres, R. Muscle Contraction Induces Acute Hydroxymethylation of the Exercise-Responsive Gene Nr4a3. Front Endocrinol (Lausanne) 7, 165, doi:10.3389/fendo.2016.00165 (2016).

58 Smemo, S. et al. Obesity-associated variants within FTO form long-range functional connections with IRX3. Nature 507, 371–375, doi:10.1038/nature13138 (2014).

59 Khokha, M. K., Hsu, D., Brunet, L. J., Dionne, M. S. & Harland, R. M. Gremlin is the BMP antagonist required for maintenance of Shh and Fgf signals during limb patterning. Nat Genet 34, 303–307, doi:10.1038/ng1178 (2003).

60 Zuniga, A., Haramis, A. P., McMahon, A. P. & Zeller, R. Signal relay by BMP antagonism controls the SHH/FGF4 feedback loop in vertebrate limb buds. Nature 401, 598–602, doi:10.1038/44157 (1999).

61 Davis, H. et al. Aberrant epithelial GREM1 expression initiates colonic tumorigenesis from cells outside the stem cell niche. Nat Med 21, 62–70, doi:10.1038/nm.3750 (2015).

62 Verheyden, J. M. & Sun, X. An Fgf/Gremlin inhibitory feedback loop triggers termination of limb bud outgrowth. Nature 454, 638–641, doi:10.1038/nature07085 (2008).

63 Hedjazifar, S. et al. The Novel Adipokine Gremlin 1 Antagonizes Insulin Action and is Increased in Type 2 Diabetes and NAFLD/NASH. Diabetes, doi:10.2337/db19-0701 (2019).

64 Bischoff, R. Regeneration of single skeletal muscle fibers in vitro. Anat Rec 182, 215–235, doi:10.1002/ar.1091820207 (1975).

65 Megeney, L. A., Kablar, B., Garrett, K., Anderson, J. E. & Rudnicki, M. A. MyoD is required for myogenic stem cell function in adult skeletal muscle. Genes Dev 10, 1173–1183, doi:10.1101/gad.10.10.1173 (1996).

66 He, S., Nakada, D. & Morrison, S. J. Mechanisms of stem cell self-renewal. Annu Rev Cell Dev Biol 25, 377–406, doi:10.1146/annurev.cellbio.042308.113248 (2009).

67 Collins, C. A. et al. Stem cell function, self-renewal, and behavioral heterogeneity of cells from the adult muscle satellite cell niche. Cell 122, 289–301, doi:10.1016/j.cell.2005.05.010 (2005).

68 Kitamoto, T. & Hanaoka, K. Notch3 null mutation in mice causes muscle hyperplasia by repetitive muscle regeneration. Stem Cells 28, 2205–2216, doi:10.1002/stem.547 (2010).

69 Lund, J. et al. Exercise in vivo marks human myotubes in vitro: Training-induced increase in lipid metabolism. PLoS One 12, e0175441, doi:10.1371/journal.pone.0175441 (2017).

70 Bergstrom, J. Percutaneous needle biopsy of skeletal muscle in physiological and clinical research. Scand J Clin Lab Invest 35, 609–616 (1975).

71 Laurens, C. et al. Perilipin 5 fine-tunes lipid oxidation to metabolic demand and protects against lipotoxicity in skeletal muscle. Sci Rep 6, 38310, doi:10.1038/srep38310 (2016).

72 Nylander, V. et al. Ionizing Radiation Potentiates High-Fat Diet-Induced Insulin Resistance and Reprograms Skeletal Muscle and Adipose Progenitor Cells. Diabetes 65, 3573–3584, doi:10.2337/db16-0364 (2016).

73 Liao, Y., Smyth, G. K. & Shi, W. The Subread aligner: fast, accurate and scalable read mapping by seed-and-vote. Nucleic Acids Res 41, e108, doi:10.1093/nar/gkt214 (2013).

74 Li, Q., Brown, J. B., Huang, H. & Bickel, P. J. Measuring reproducibility of high-throughput experiments. The Annals of Applied Statistics 5, 1752–1779 (2011).

75 Liao, Y., Smyth, G. K. & Shi, W. featureCounts: an efficient general purpose program for assigning sequence reads to genomic features. Bioinformatics 30, 923–930, doi:10.1093/bioinformatics/btt656 (2014).

76 Robinson, M. D., McCarthy, D. J. & Smyth, G. K. edgeR: a Bioconductor package for differential expression analysis of digital gene expression data. Bioinformatics 26, 139–140, doi:10.1093/bioinformatics/btp616 (2010).

77 Lun, A. T. & Smyth, G. K. De novo detection of differentially bound regions for ChIP-seq data using peaks and windows: controlling error rates correctly. Nucleic Acids Res 42, e95, doi:10.1093/nar/gku351 (2014).

78 Yu, G., Wang, L. G. & He, Q. Y. ChIPseeker: an R/Bioconductor package for ChIP peak annotation, comparison and visualization. Bioinformatics 31, 2382–2383, doi:10.1093/bioinformatics/btv145 (2015).

79 RCoreTeam. R: A language and environment for statistical computing. R Foundation for Statistical Computing, Vienna, Austria. (2018).

80 Douglas, B., Martin, M., Ben, B. & Steve, W. Fitting Linear Mixed-Effects Models Using lme4. Journal of Statistical Software 67, 1–48, doi:10.18637 (2015).

81 Lenth, R. emmeans: Estimated Marginal Means, aka Least-Squares Means. R package version 1.3.4. (2019).

82 Wickham, H. ggplot2: Elegant Graphics for Data Analysis. Springer-Verlag New York (2016).

83 Ji, H. et al. An integrated software system for analyzing ChIP-chip and ChIP-seq data. Nat Biotechnol 26, 1293–1300, doi:10.1038/nbt.1505 (2008).

84 Chakroun, I. et al. Genome-wide association between Six4, MyoD, and the histone demethylase Utx during myogenesis. FASEB J 29, 4738–4755, doi:10.1096/fj.15-277053 (2015).

85 Pfaffl, M. W. A new mathematical model for relative quantification in real-time RT-PCR. Nucleic Acids Res 29, e45, doi:10.1093/nar/29.9.e45 (2001).

86 Rappsilber, J., Ishihama, Y. & Mann, M. Stop and go extraction tips for matrix-assisted laser desorption/ionization, nanoelectrospray, and LC/MS sample pretreatment in proteomics. Anal Chem 75, 663–670, doi:10.1021/ac026117i (2003).

87 Scheltema, R. A. et al. The Q Exactive HF, a Benchtop mass spectrometer with a pre-filter, high-performance quadrupole and an ultra-high-field Orbitrap analyzer. Mol Cell Proteomics 13, 3698–3708, doi:10.1074/mcp.M114.043489 (2014).

88 Cox, J. & Mann, M. MaxQuant enables high peptide identification rates, individualized p.p.b.-range mass accuracies and proteome-wide protein quantification. Nat Biotechnol 26, 1367–1372, doi:10.1038/nbt.1511 (2008).

89 Cox, J. et al. Accurate proteome-wide label-free quantification by delayed normalization and maximal peptide ratio extraction, termed MaxLFQ. Mol Cell Proteomics 13, 2513–2526, doi:10.1074/mcp.M113.031591 (2014).

90 Pirkmajer, S. et al. Methotrexate promotes glucose uptake and lipid oxidation in skeletal muscle via AMPK activation. Diabetes 64, 360–369, doi:10.2337/db14-0508 (2015).

91 Mounier, R. et al. Important role for AMPKalpha1 in limiting skeletal muscle cell hypertrophy. FASEB J 23, 2264–2273, doi:10.1096/fj.08-119057 (2009).

